# Characterization of sequence specific binding of LARP6 to the 5’ stem-loop of type I collagen mRNAs and implications for rational design of antifibrotic drugs

**DOI:** 10.1101/2020.09.30.320564

**Authors:** Lela Stefanovic, Blaine H. Gordon, Robert Silvers, Branko Stefanovic

## Abstract

Excessive synthesis of type I collagen characterizes fibrotic diseases. Binding of LARP6 to the 5’ stem-loop (5’SL) of collagen mRNAs regulates their translation and the high rate of biosynthesis in fibrosis. LARP6 needs two domains to form stable complex with 5’SL RNA, the La-domain and the juxtaposed RRM domain (jointly called the La-module). We describe that the La-domain of LARP6 is necessary and sufficient for recognition of 5’SL in sequence specific manner. The three amino acid motif, RNK, located in the flexible loop which connects the second α-helix to the β-sheet of the La domain is critical for binding. Mutation of any of these three amino acids abolishes the binding of La-domain to 5’SL. The major site of crosslinking of LARP6 to 5’SL RNA was mapped to this motif. The RNK motif is not found in other LARPs, which can not bind 5’SL. Presence of RRM increases the stability of complex between La-domain and 5’SL RNA and RRM domain does not make extensive contacts with 5’SL RNA. We propose a model in which the initial recognition of 5’SL by LARP6 is mediated by the RNK epitope and further stabilized by the RRM domain. This discovery suggests that the interaction between LARP6 and collagen mRNAs can be blocked by small molecules that target the RNK epitope and will help rational design of the LARP6 binding inhibitors as specific antifibrotic drugs.

## Introduction

LARP6 (La ribonucleoprotein domain family member 6) is a member of LARP superfamily of RNA binding proteins, which are regulators of translation [1–5]. While other members of the superfamily bind either homopolymers of RNA or show no strict sequence preference [6–8], LARP6 binds a unique sequence. This sequence is found in two mRNAs encoding for collagen α1(I) and α2(I) polypeptides, which fold into type I collagen, and in collagen α1(III) mRNA which encodes for type III collagen [9]. This sequence was termed collagen 5’ stem-loop (5’SL), and is located in the 5’UTR of these mRNAs overlapping the start codon. The 5’SL is 48 nucleotides long and folds into two stems flanking a central bulge of 9 nucleotides. Our previous work has identified 5 nucleotides within the bulge which are critical for binding LARP6; mutation of any of these 5 nucleotides abolished the interaction of 5’SL with LARP6 in gel mobility shift experiments [10]. The stems are important only as dsRNA that flanks the bulge.

It has been reported that two domains of LARP6, the La domain and the RRM domain (together called the La module) are necessary for interaction with the 5’SL [11]. The involvement of two protein domains and five nucleotides of RNA suggested complex interactions between the 5’SL and LARP6. The Kd of 5’SL/LARP6 interaction was measured to be 0.4 nM using fluorescence polarization [10], 12 nM using gel mobility shift assay [12] and 40 nM using ITC [11].

Collagens type I and III are fibrilar proteins expressed in bone, tendons and skin, with type I representing 90% of fibrilar collagen in these organs [13]. In the pathological state of fibrosis type I collagen is expressed at high levels in other organs, such as liver, heart, kidney, blood vessels, joints or lungs, leading to their scarring and loss of function. Primary and secondary fibrosis affect 45% of population [14] and represent a major medical problem, however, there is no cure for fibrosis. Type I collagen is composed of two α1(I) and one α2(I) polypeptides and its formation is highly dependent on local concentration of the individual polypeptides, because all three polypeptides have to register their C-terminal domains to initiate the nucleation of heterotrimer [15]. In normal tissues type I collagen is one of the most stable proteins with half life of 4-12 months and its fractional synthesis rate (FSR, defined as % synthesis per day) is about 2% in the skin [16], while in the liver it is only 0.2% [17]. This is significantly slower than the FSR of most proteins (18-140% per day) [18]. In fibrosis, however, the synthesis of type I collagen is dramatically increased [19, 20]. Binding of LARP6 to collagen mRNAs is necessary for this increase [21], because LARP6 recruits accessory translational factors to increase translational competency of collagen mRNAs and to couple translation of α1(I) polypeptide to that of α2(I) polypeptide [2, 22–26]. The coupled translation of collagen mRNAs consolidates production of both polypeptides at discrete sites on the endoplasmic reticulum (ER) membrane; this increases the local concentration of polypeptides and accelerates their folding into type I procollagen.

The experimental animals which synthesize type I collagen without LARP6 regulation are resistant to hepatic fibrosis. [27], indicating the importance of LARP6 dependent regulation in fibrosis. This resulted in a concept that inhibitors of LARP6 binding to 5’SL can be specific antifibrotic drugs. We have discovered and characterized one such inhibitor by high throughput screening [28] and developed a phenotypic screening method for additional drug screening efforts [29]. Here we describe the molecular characterization of sequence specific interaction of LARP6 and 5’SL RNA which will help rational design of additional inhibitors.

## Results

### La-domain of human LARP6 is sufficient for sequence specific binding of 5’SL RNA

Fig 1A shows schematic representation of the domains of human LARP6. Two domains, the La and RRM domains, are jointly called the La-module [4]. La-module, as well as full size LARP6, binds α1(I) 5’SL and α2(I) 5’SL with similar affinity (the sequences of these elements is shown in Fig 1B) [10]. In this work we have used both 5’SL RNA sequences, as indicated. It has been reported before that the entire La-module is necessary for binding α1(I) 5’SL RNA [11]. However, using recombinant proteins or cell extracts immediately after preparation, we were able to show that La-domain alone is sufficient to bind 5’SL in the sequence specific manner.

**Figure 1.**
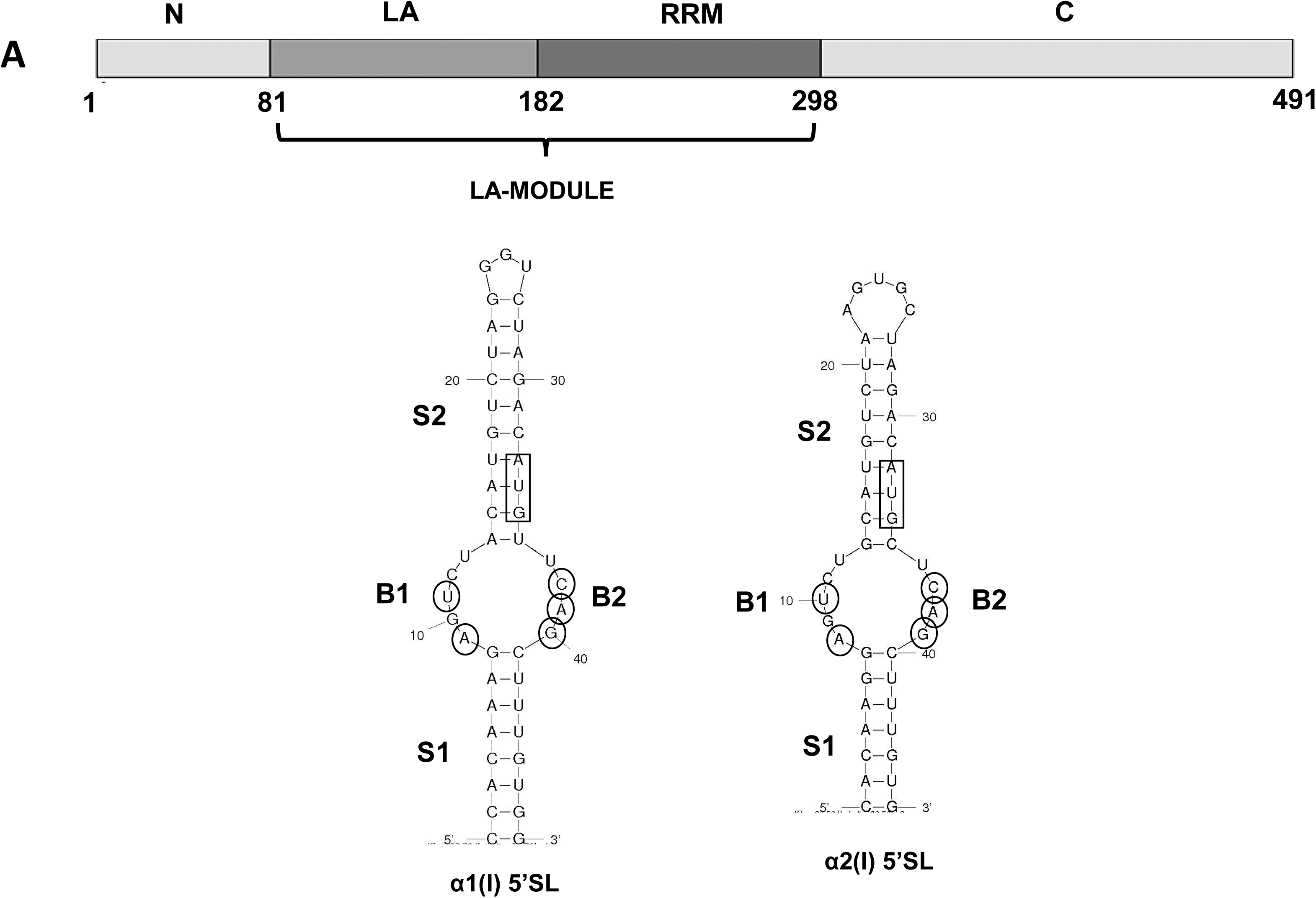
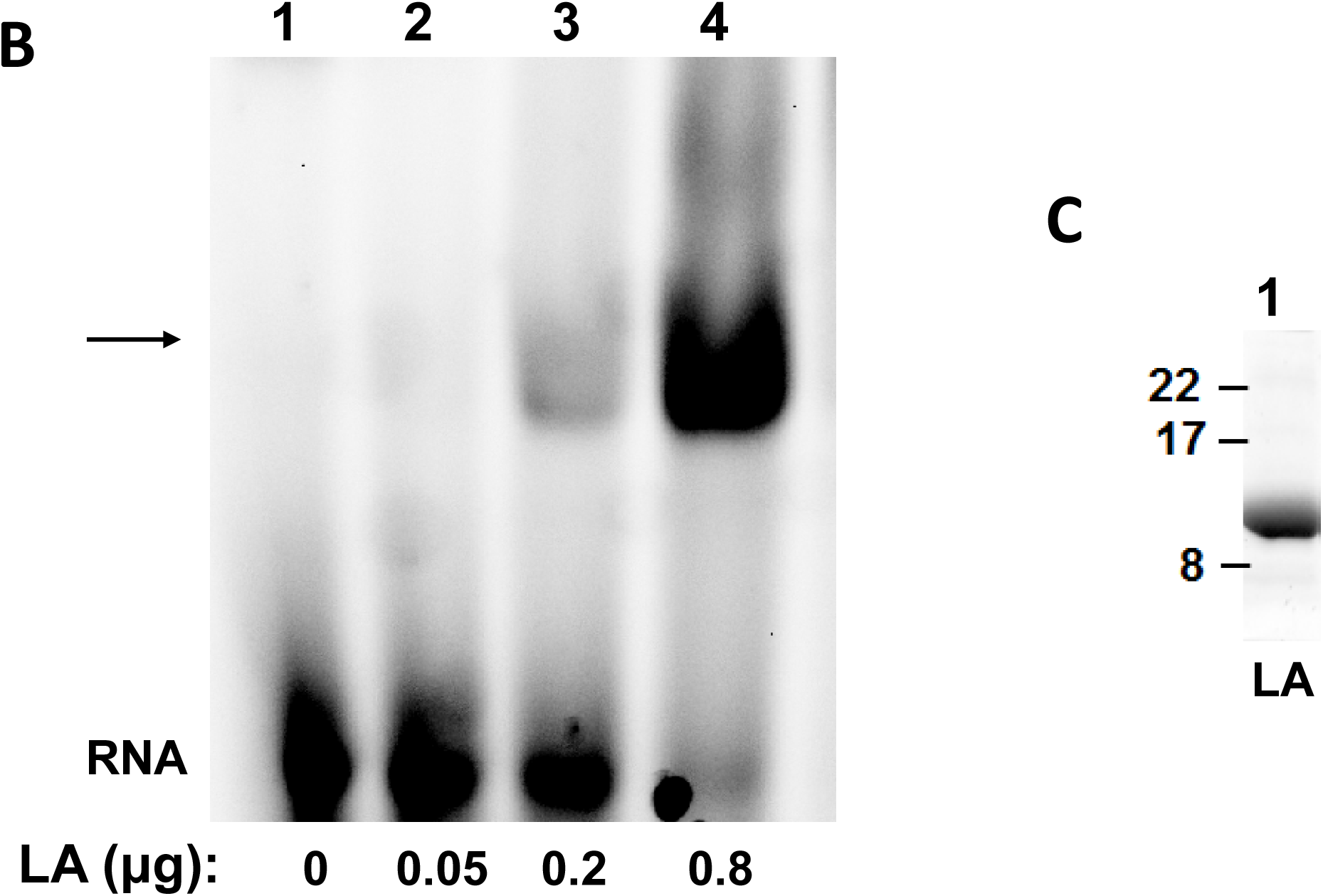
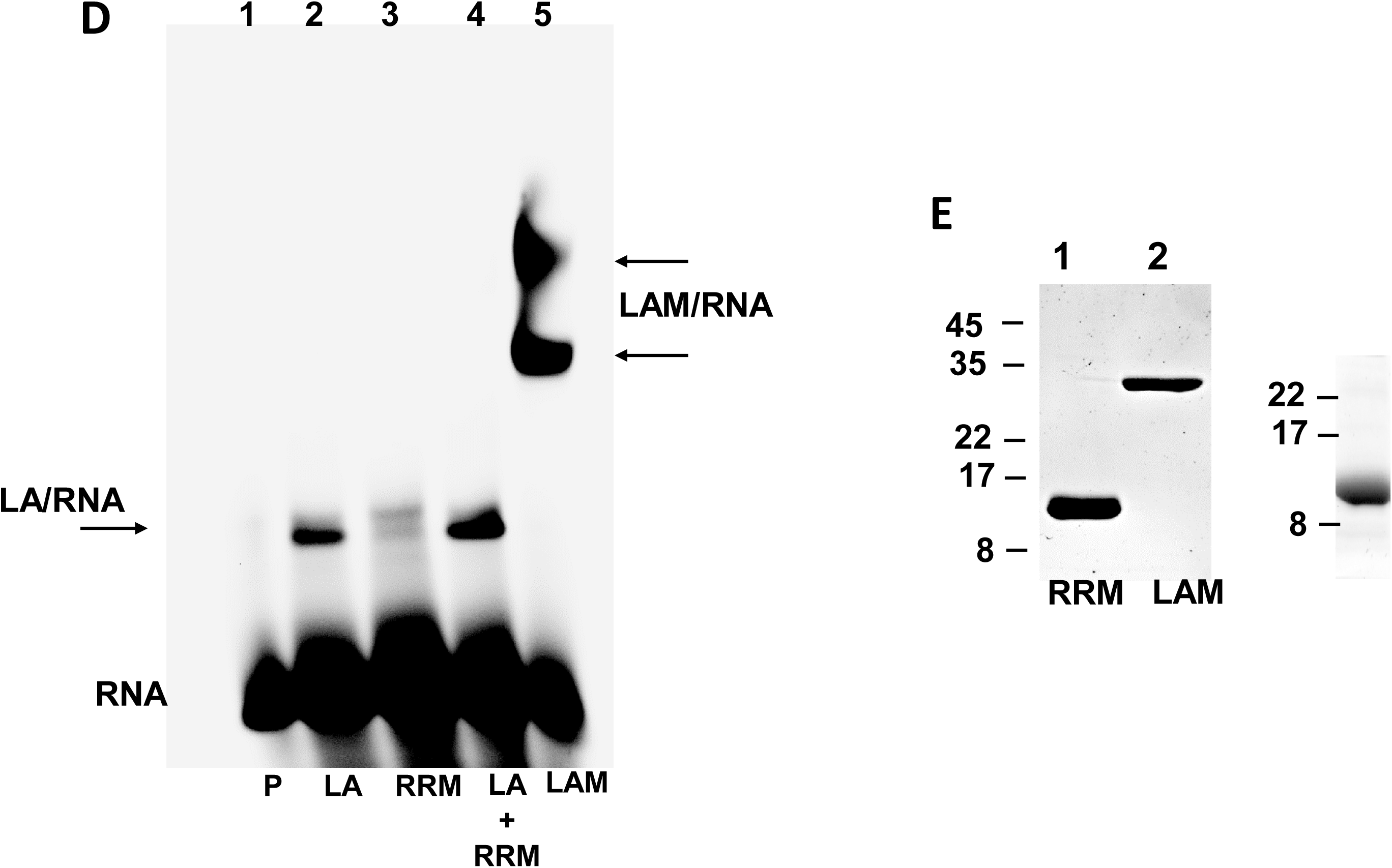
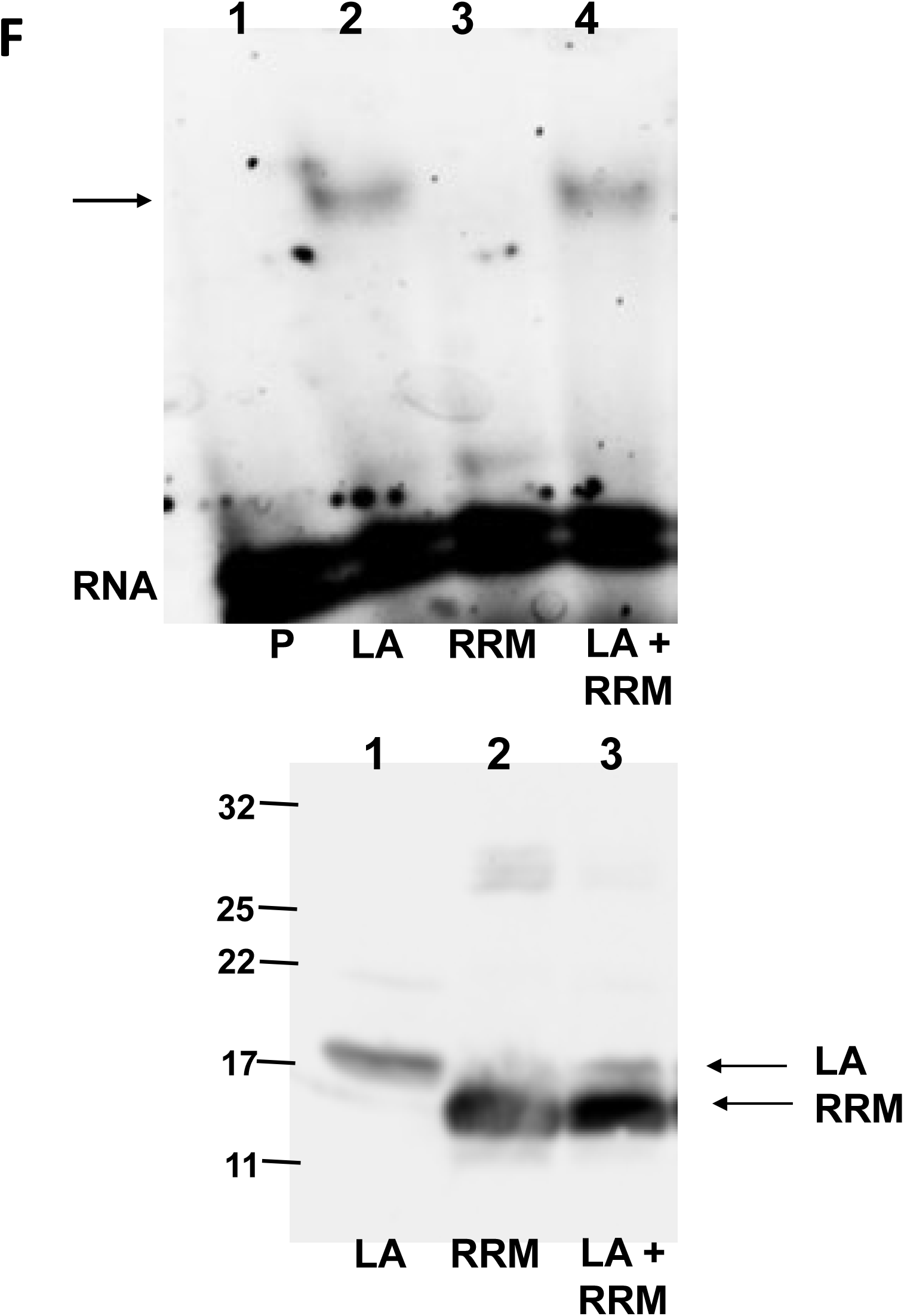
La-domain of LARP6 binds 5’SL RNA. A. Schematic representation of the domains of LARP6. La-module is indicated. LA, La-domain, RRM, RNA recognition domain, N and C, N-terminal and C-terminal domains. Sequences of human collagen 5’SL of collagen α1(I) and α2(I) mRNAs are shown below. The nucleotides important for binding LARP6 are circled. B1 and B2, single stranded parts of the central bulge. S1 and S2, stems. Translational initiation codon is boxed. B. Binding of recombinant La-domain to α2(I) 5’SL RNA. Purified La domain was added at increasing amounts to α2(I) 5’SL RNA probe (RNA) and complexes were resolved by gel mobility shift assay. Arrow indicates migration of La/5’SL complex. C. Recombinant La domain visualized by Coomassie staining. D. Binding of recombinant RRM domain to α2(I) 5’SL RNA. Recombinant La domain (lane 2), recombinant RRM domain (lane 3), an equimolar mix of La and RRM domains (lane 4) or recombinant LaM (lane 5) were added to α2(I) 5’SL RNA and complexes (arrows) resolved by gel mobility shift assay. Lane 1, 5’SL RNA alone (P). E. Recombinant RRM and LAM visualized by Coomassie staining. F. Binding of La domain expressed in mammalian cells. Upper panel: gel mobility shift with α2(I) 5’SL RNA probe and extracts of cells expressing La-domain (lane 2, LA) or RRM (lane 3, RRM) or co-expressing both domains in trans (lane 4). Lane 1 is RNA probe alone. Arrow indicates La/5’SL complex. Bottom panel: expression of transfected proteins in the same extract.

First, we expressed recombinant La-domain (AA 81-182) in E. coli and purified the protein by single step purification (Fig 1C). We observed that excessive purifications, as well as freezing and thawing of the protein, inactivates its binding activity. Therefore, we used only fresh preparations in the course of this work. If present, the contamination by bacterial proteins has no effect on binding of La domain to 5’SL RNA (supplementary figure 1). Fig 1B shows that when recombinant La-domain was added in increasing amounts to α2(I) 5’SL RNA, RNA/protein complex was formed, which was stable enough to withstand gel electrophoresis. We also expressed RRM-domain (AA 183-298) in bacteria and analyzed its binding to α2(I) 5’SL (Fig 1D). While La-domain showed binding activity to 5’SL RNA (lane 2), RRM-domain was unable to form a complex (lane 3). When RRM-domain was mixed with La-domain the binding was indistinguishable from that of La-domain alone (compare lanes 4 and 2). This suggested that RRM can not complement the binding of La-domain when present in trans. As control, we also analyzed the binding of recombinant La-module (LAM, lane 5), where both domains are in cis. La-module showed strong binding with formation of two complexes; these complexes represent monomer and dimer of LAM bound to RNA, what has been occasionally seen with LAM preparations. The RRM and LAM proteins are shown in Fig 1E.

To verify these results with the La-domain expressed in mammalian cells we cloned individually the HA-tagged La-domain and RRM domain into mammalian expression vector, expressed the domains in HEK293 cells and analyzed their binding activity to α2(I) 5’SL RNA in whole cell extracts by gel mobility shift assay. We found that the La-domain alone was able to interact with 5’SL RNA probe (Fig 1F, lane 2), while the RRM domain was not (lane 3). When both domains were co-expressed in trans, the binding was indistinguishable to that of the La-domain alone (lane 4). Expression level of the domains was analyzed by western blot using anti-HA antibody (Fig 1F, bottom panel). The RRM domain was expressed at higher level than the La-domain, but no binding activity was evident. This suggested that the La-domain is necessary and sufficient for recognition of 5’SL in mammalian cells and that RRM-domain alone has no binding activity towards 5’SL RNA.

La-domain alone accumulated equally in the nucleus and in the cytoplasm of cells, as assessed by immunostaining (supplementary Fig 2). Full size LARP6 is predominantly cytoplasmic and has distinct peri-nuclear accumulation, not seen with the La-domain alone. We have reported previously that LARP6 associates with the ER membrane [30] and the peri-nuclear localization is consistent with its role in translation of collagen mRNAs on the ER membrane.

**Figure 2.**
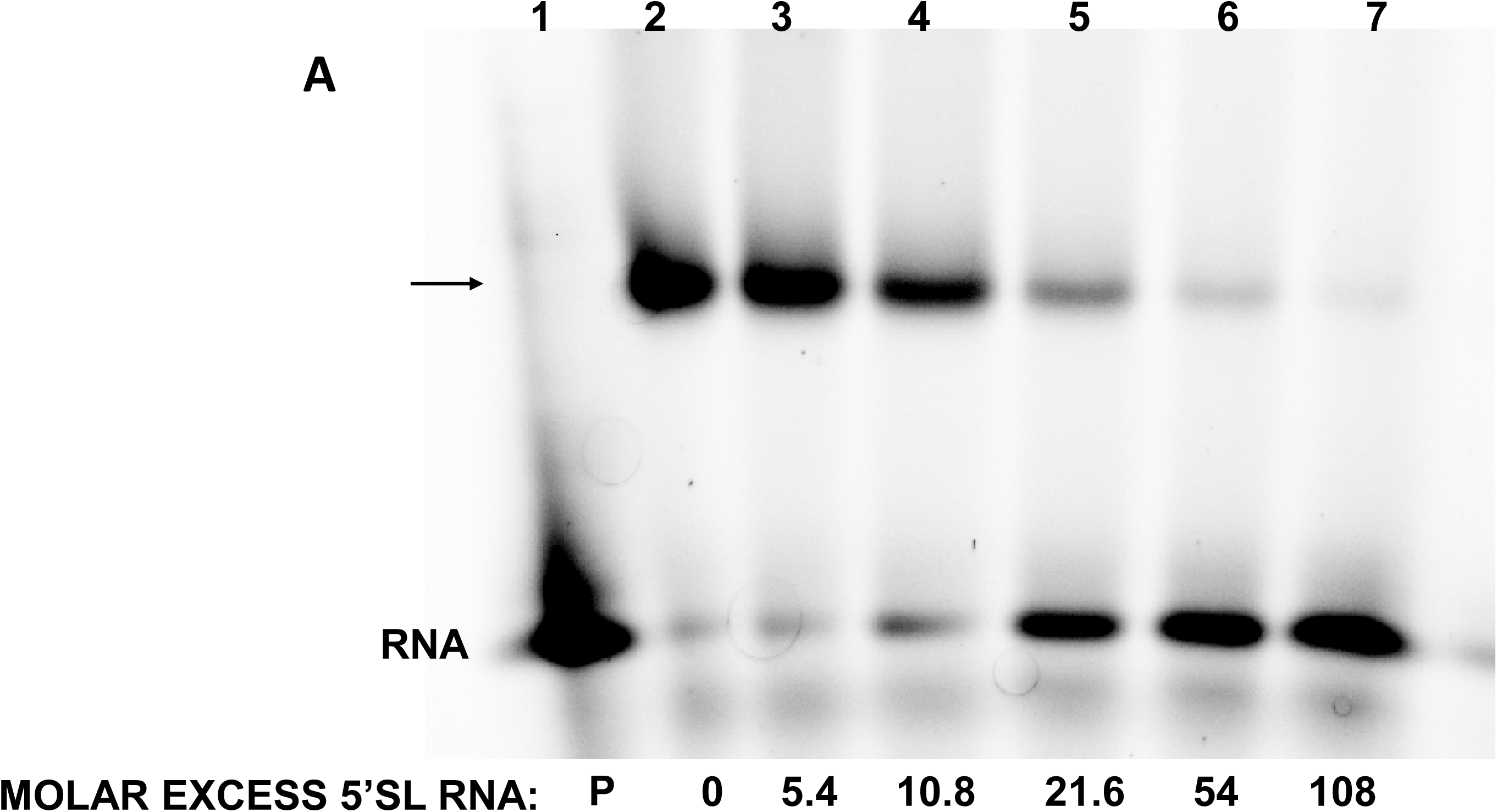
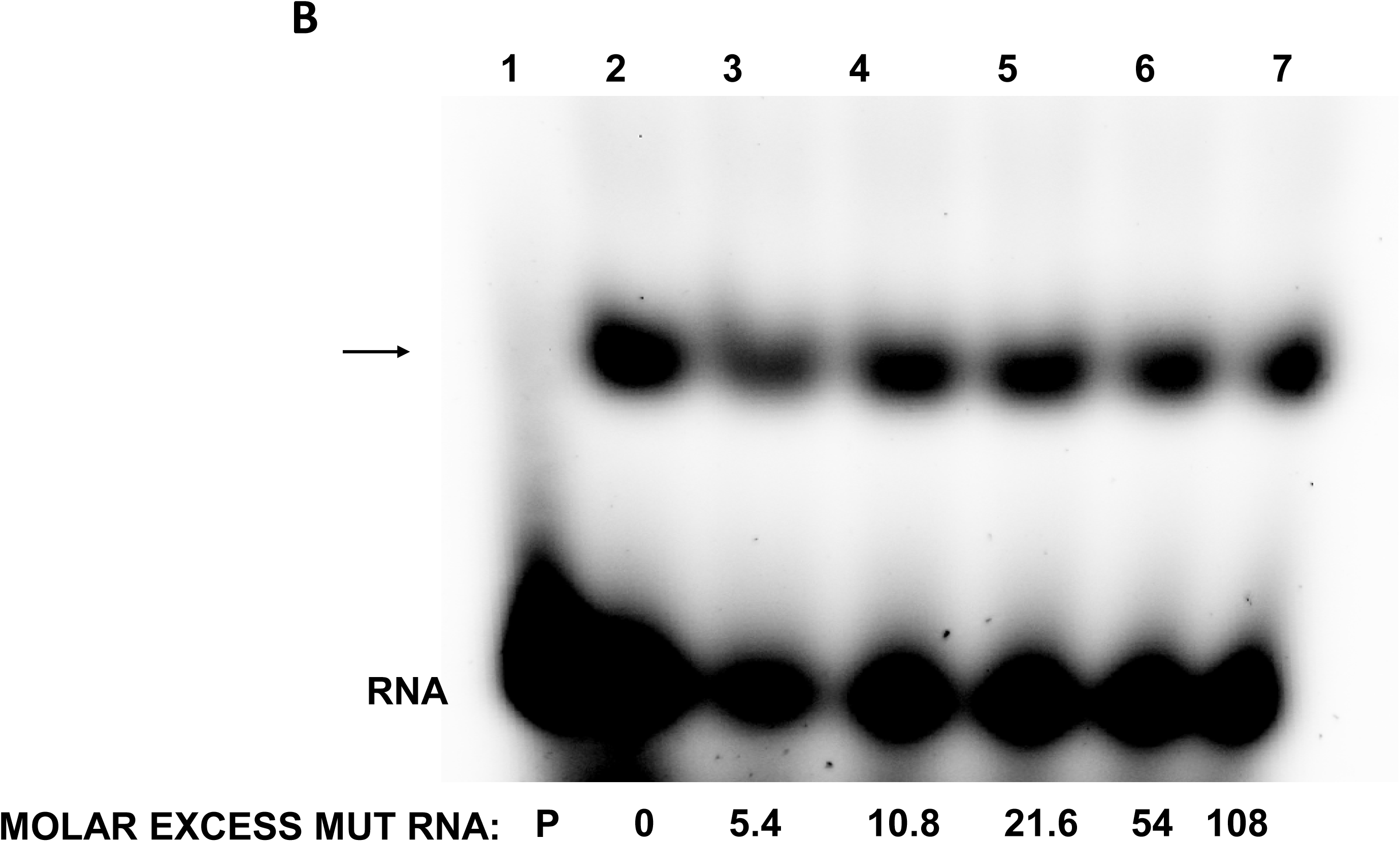
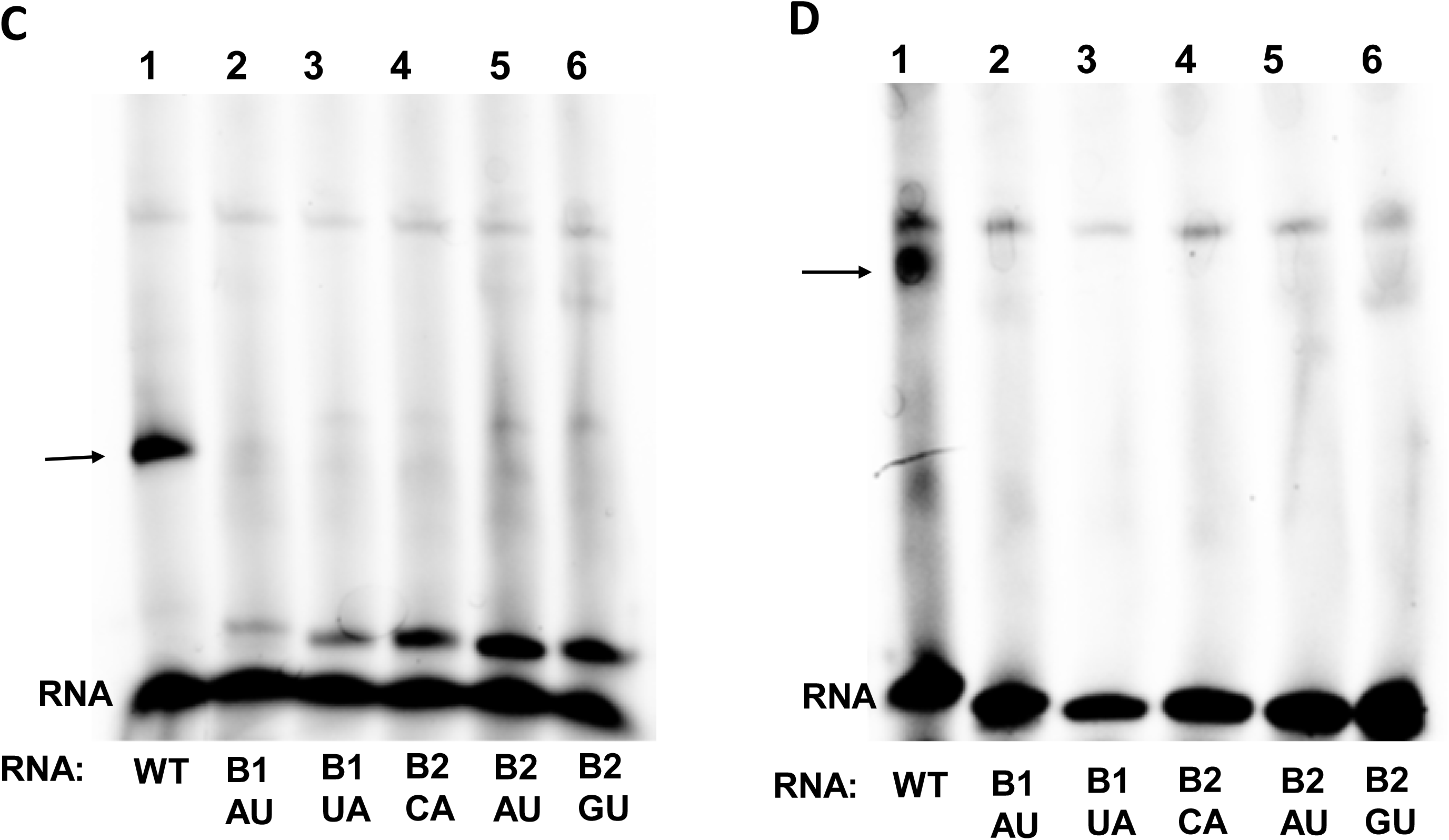
Sequence specific binding of La-domain to 5’SL RNA. A. Competition of La-domain binding by specific competitor RNA. Binding of recombinant La-domain to α2(I) 5’SL RNA probe was analyzed in the presence of indicated molar excess of unlabeled α2(I) 5’SL RNA by gel mobility shift. P, probe alone (lane 1). Arrow indicates La/5’SL complex. B. Competition by mutant RNA. Same experiment as in D, except molar excess of mutant 5’SL RNA was used. C. Binding of recombinant La-domain to the mutants of 5’SL. WT α1(I) 5’SL RNA probe (lane 1) or mutant probes in which individual nucleotides of the bulge had been mutated (lanes 2-6) were incubated with recombinant La-domain and complexes analyzed by gel mobility shift. Arrow indicates La/5’SL complex. D. Experiment as in C, except recombinant La-module was used. Arrow indicates LaM/5’SL complex.

To analyze if binding of La domain to 5’SL is sequence specific we employed competition experiments and used mutant 5’SL RNA probes. First, we competed the binding of recombinant La-domain to labeled α1(I) 5’SL RNA with excess of unlabeled wt α1(I) 5’SL RNA (Fig 2A). After 30 min of equilibration we analyzed the complex on labeled 5’SL RNA by gel mobility shift. In the presence of wt 5’SL competitor the complex started to diminish at ~10-fold molar excess of the competitor (Fig 2A, lane 4) and almost complete dissociation was seen with 100-fold excess of the competitor (lane 7). Next, we competed the complexes with unlabeled mutant 5’SL RNA in which we substituted the five nucleotides critical for binding of LARP6. The mutant 5’SL competitor had no effect on the La/5’SL complex even at >100-fold molar excess (Fig 1B), suggesting the sequence specific binding.

The 5 nucleotides of α1(I) 5’SL which are critical for binding LARP6 are indicated in Fig 1A [2, 10]. Mutation of each of these nucleotides abolished the binding of LARP6 or La-module to 5’SL. Now we assessed if La-domain alone can distinguish between the single nucleotide mutants of α1(I) 5’SL. We used the 5’SL probes in which each of the critical nucleotides (circled in Fig 1) was mutated and analyzed binding of recombinant La-domain by gel mobility shift assay. Fig 2C shows that only wt α1(I) 5’SL RNA was recognized by the La-domain (lane 1), while all mutant α1(I) 5’SL RNAs failed to bind. For comparison, Fig 2D shows the specificity of binding of recombinant La-module; La-module also recognized only the wt α1(I) 5’SL RNA (lane 1). Thus, the recombinant La-domain binds α1(I) 5’SL RNA with the same sequence specificity as the entire La-module, suggesting that the La-domain contains all the structural information needed for the sequence specific recognition of 5’SL.

### RRM domain increases the stability of binding to 5’SL RNA

We have previously described a fluorescence polarization (FP) method to assess the formation of LARP6/5’SL complexes. When increasing amounts of fluorescently labeled 5’SL RNA is added to a fixed amount of recombinant proteins, the RNA binds only to the active protein molecules. The formation of RNA/protein complexes as a function of the RNA concentration is monitored as increase in FP and the affinity of binding in different protein preparations can be compared. Such experiment with α2(I) 5’SL RNA and recombinant La-domain and La-module is shown in Fig 3A. The α2(I) 5’SL RNA bound La-domain and La-module with similar kinetics, suggesting that both proteins are recognized with similar affinity.

**Figure 3.**
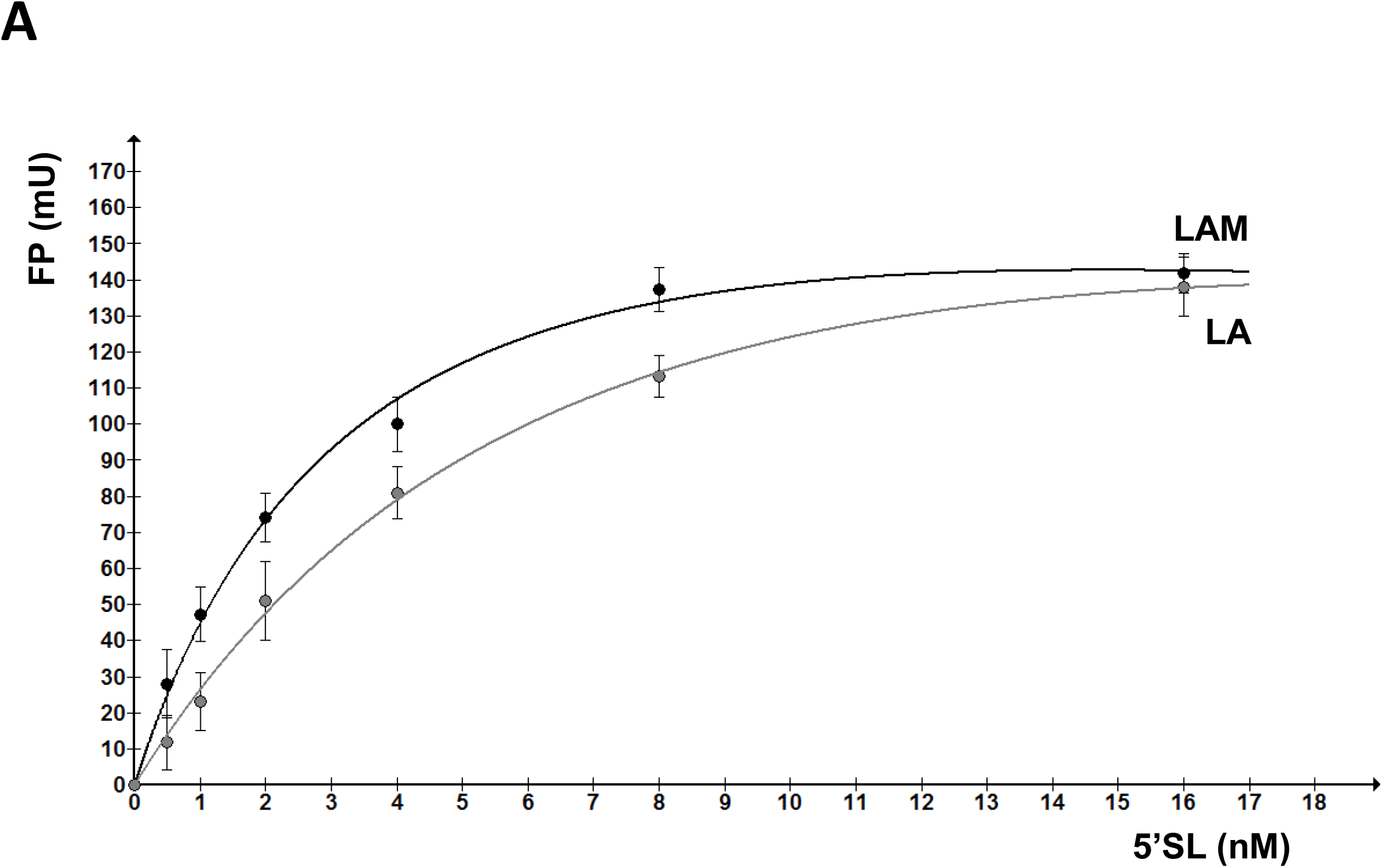
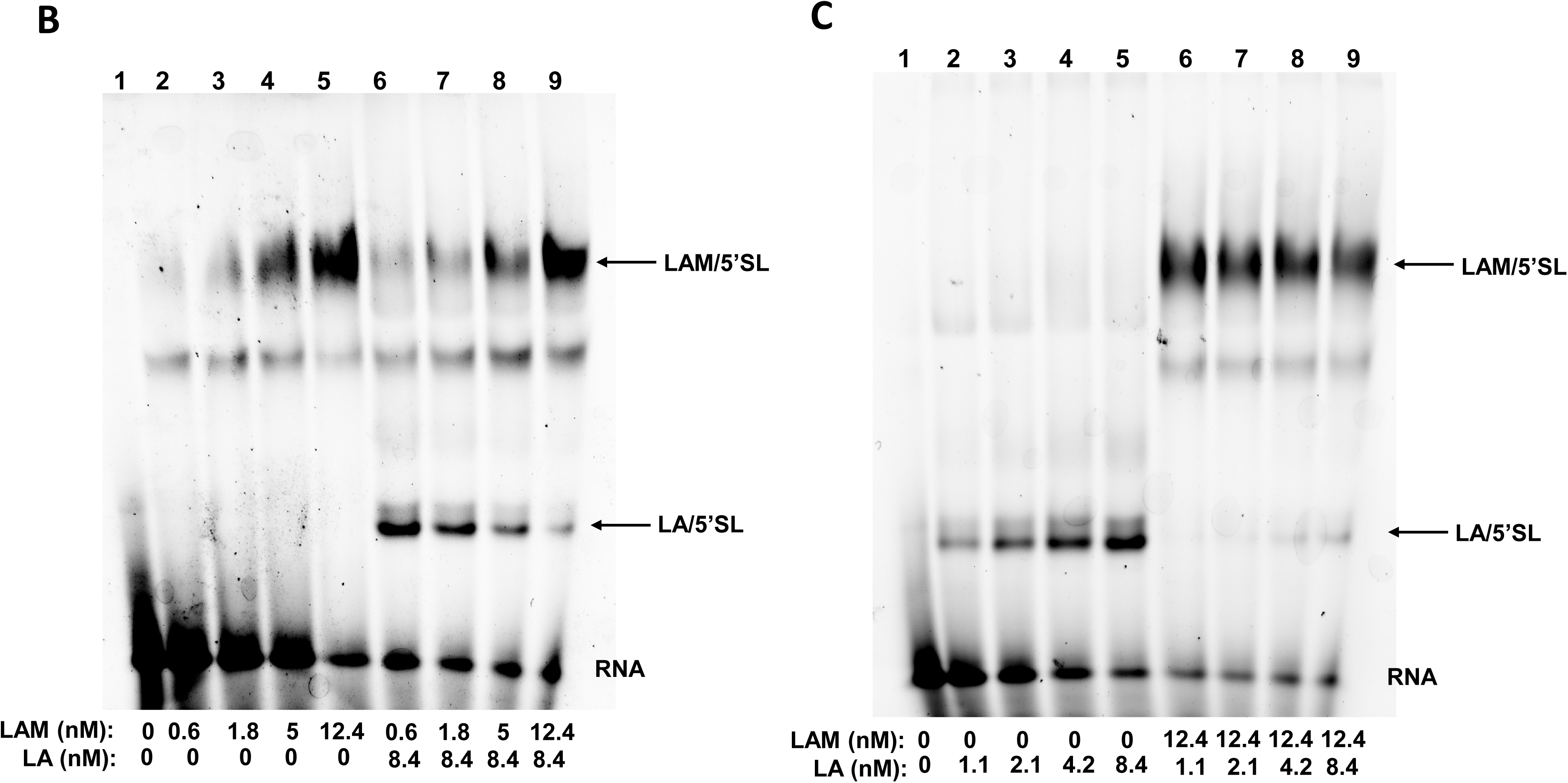
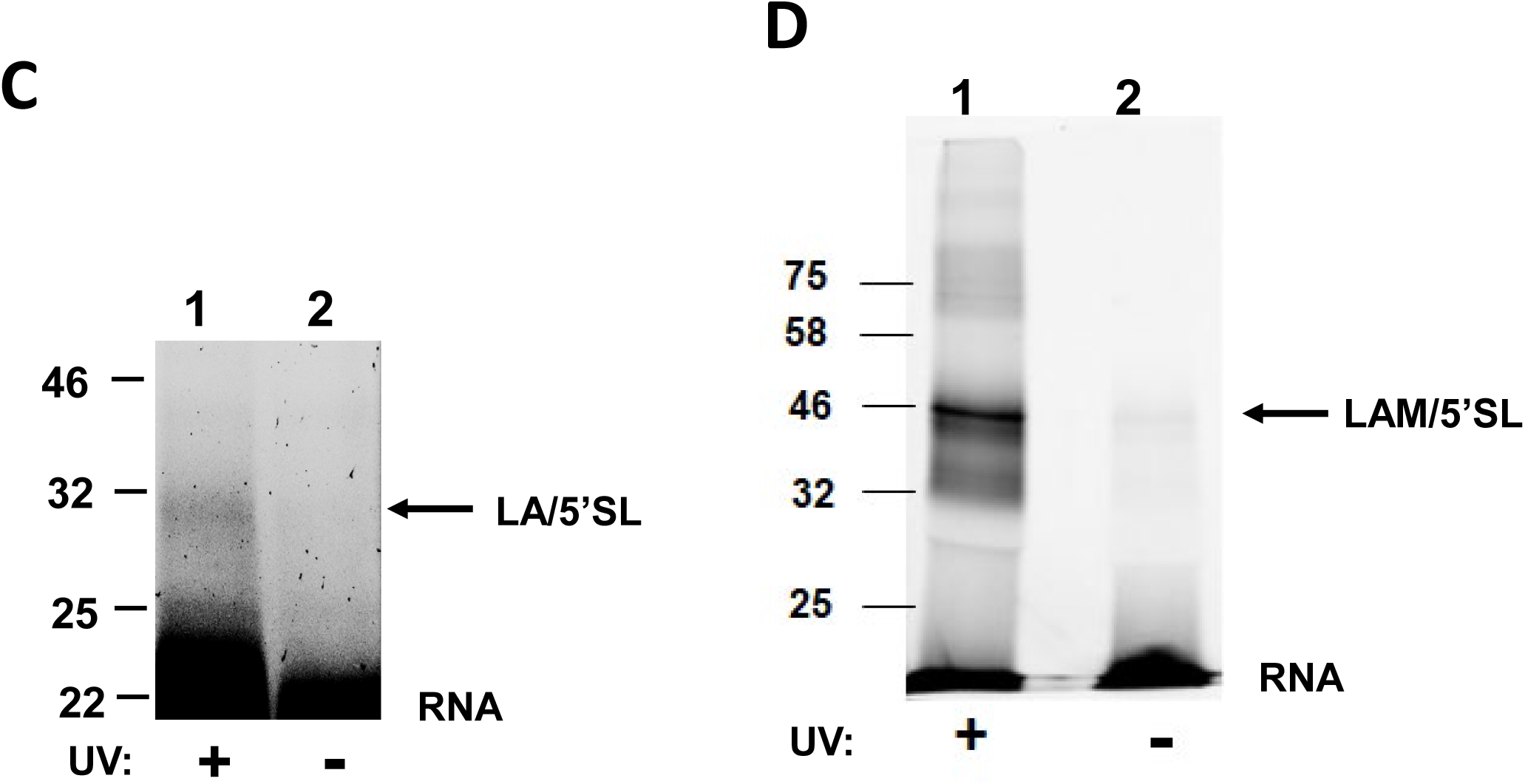
Cross-competition of La-domain and La-module for 5’SL RNA. A. Titration of La-domain and La-module with increasing amounts of α2(I) 5’SL RNA. Increasing amounts of fluorescently labeled α2(I) 5’SL RNA were added to La-domain (black line) or La-module (gray line) at 3.8 μM and fluorescence polarization (FP) was measured and plotted against 5’SL RNA concentration. N=6, error bars: +/− 1SD. B. Competition of La-module binding by La-domain. Increasing amounts of recombinant La-module were incubated with α2(I) 5’SL RNA probe in absence (lanes 2-5) or in presence (lanes 6-9) of fixed amount of recombinant La-domain and complexes were resolved by gel electrophoresis. Lane 1, probe alone. Arrows indicate migration of the respective complexes. C. Competition of La-domain binding by La-module. Experiment as in A, except increasing amounts of recombinant La-domain were incubated with α2(I) 5’SL RNA probe in absence (lanes 2-5) or in presence (lanes 6-9) of fixed amount of recombinant La-module. C. UV crosslinking of La-domain and La-module to 5’SL RNA. Recombinant La-domain (left panel) and La-module (right panel) were bound to α2(I) 5’SL RNA probe and separated on denaturing SDS-PAGE gel after UV crosslinking (lane 1) or without UV crosslinking (lane 2) and visualized by imaging of RNA fluorescence. Arrows indicate protein/RNA complex.

Next, we tested the stability of La/5’SL and LAM/5’SL complexes by performing cross-competition experiments. The difference in electrophoretic mobility between 5’SL RNA in complex with La-domain and in complex with La-module allowed us to analyze both complexes simultaneously when the proteins compete for 5’SL RNA. In one experiment (Fig 3B) we prepared increasing amounts of recombinant La-module in absence (lanes 2-5) or presence (lanes 6-9) of fixed amount of La-domain, added labeled 5’SL RNA and analyzed the equilibration of complexes by gel mobility assay. The increasing amounts of La-module bound increasing amounts of 5’SL RNA regardless of the absence (lanes 2-5) or presence of La-domain in the reaction (lanes 6-9), suggesting that La-domain can not compete with La-module for 5’SL RNA. At the same time, the complex formed with La-domain in presence of low amounts of La-module (lane 6) was competed out by increasing amounts of La-module (lanes 7-9).

In the opposite experiment (Fig 3B) we prepared increasing amounts of recombinant La-domain in absence (lanes 2-5) or presence (lanes 6-9) of fixed amount of La-module and added labeled 5’SL RNA. The increasing amounts of La-domain bound increasing amounts of 5’SL RNA in the absence of La-module (lanes 2-5), however, the fixed amount of La-module readily competed out the La-domain complexes (lanes 6-9). These results clearly demonstrated that combination of La/RRM (La-module) outcompetes La-domain for binding to 5’SL. This suggests that the function of RRM is to increase the stability of the complex formed with 5’SL RNA.

Crosslinking by UV light covalently attaches RNA to a protein if they are in close proximity [31]. We have previously reported a crosslink between La-module and α1(I) 5’SL RNA [2]. Here we compared the crosslinks of La-domain and La-module to α2(I) 5’SL RNA. A weak UV crosslink was formed with La-domain (Fig 3C), while a stronger crosslink was seen with La-module (Fig 3D), suggesting a more intimate contact of the La-module with 5’SL RNA.

### The length of La-domain modulates 5’SL binding

The C-terminal 15 amino-acids of La-domain are unstructured, but it has been reported that this sequence contributes to the synergistic effect of La/RRM binding [11]. To assess the role of this sequence in binding of La-domain alone we deleted amino acids from the C-terminus of La-domain, expressed the truncated domains in HEK293 cells and analyzed the binding in cell extracts by gel mobility shift (Fig 4A). Deletion of 8 or 16 amino acids at the C-terminus abolished the binding (lanes 3 and 5), suggesting that C-terminal amino acids are important. To address if this is sequence specific or length specific effect we appended random sequences to the truncated La-domain. Fig 4B, lanes 2 and 3 shows that two random sequences partially restored the binding of truncated La-domain. When we extended La-domain with 26 random amino acids the binding was increased (lane 4). From these experiments we concluded that a certain length of La domain is needed and that appending various sequences to the C-terminus can modulate the binding affinity. In this respect, RRM can be regarded as an extension of La-domain with the sequence which increases the stability La-domain/5’SL complex.

**Figure 4.**
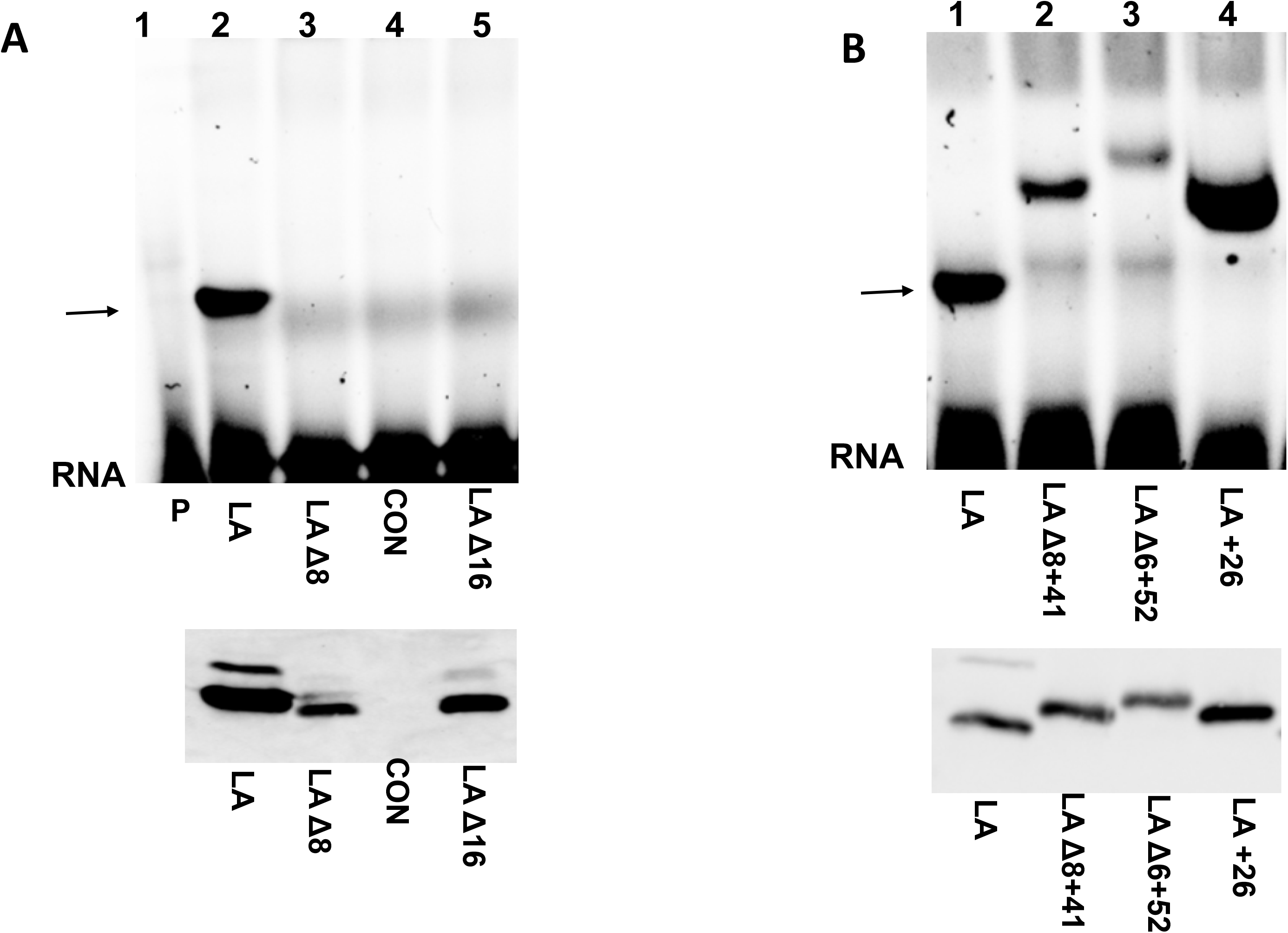
C-terminus of La-domain modulates binding. A. Effect of the shortening of C-terminus. Top panel: full size La-domain (81-182, lane 2), La-domain with eight C-terminal amino acids deleted (81-174, lane 3) or La-domain with sixteen C-terminal amino acids deleted (81-166, lane 5) were expressed in HEK293 cells and cell extract analyzed for binding to α2(I) 5’SL probe by gel mobility shift. Lane 1, probe alone, lane 4), untransfected cells. Bottom panel: expression of proteins in the same extract measured by western blot. B. Effect of extension of La-domain. Full size La-domain (81-182, lane 1) or La-domain with various random sequences appended at the C-terminus (lanes 2-4) were analyzed as in A.

### The RNK epitope of La-domain is critical for 5’SL recognition

Having established that La-domain is involved in sequence specific recognition of 5’SL, we performed site directed mutagenesis to identify the epitope in La-domain involved in 5’SL recognition. We concentrated on the three amino acids in the loop between second helix and first strand of β-sheet of La-domain [11]. These amino acids, R121, N122 and K123, are unique to LARP6 and not found in other LARPs (see below). We changed each of these amino acids into alanine, expressed the individual mutants in HEK293 cells and analyzed the binding in cell extracts by gel mobility shift. Fig 5A and 5B show that changing any of these three amino acids into alanine completely abolished the ability of La-domain to recognize α2(I) 5’SL RNA and α1(I) 5’SL.

**Figure 5.**
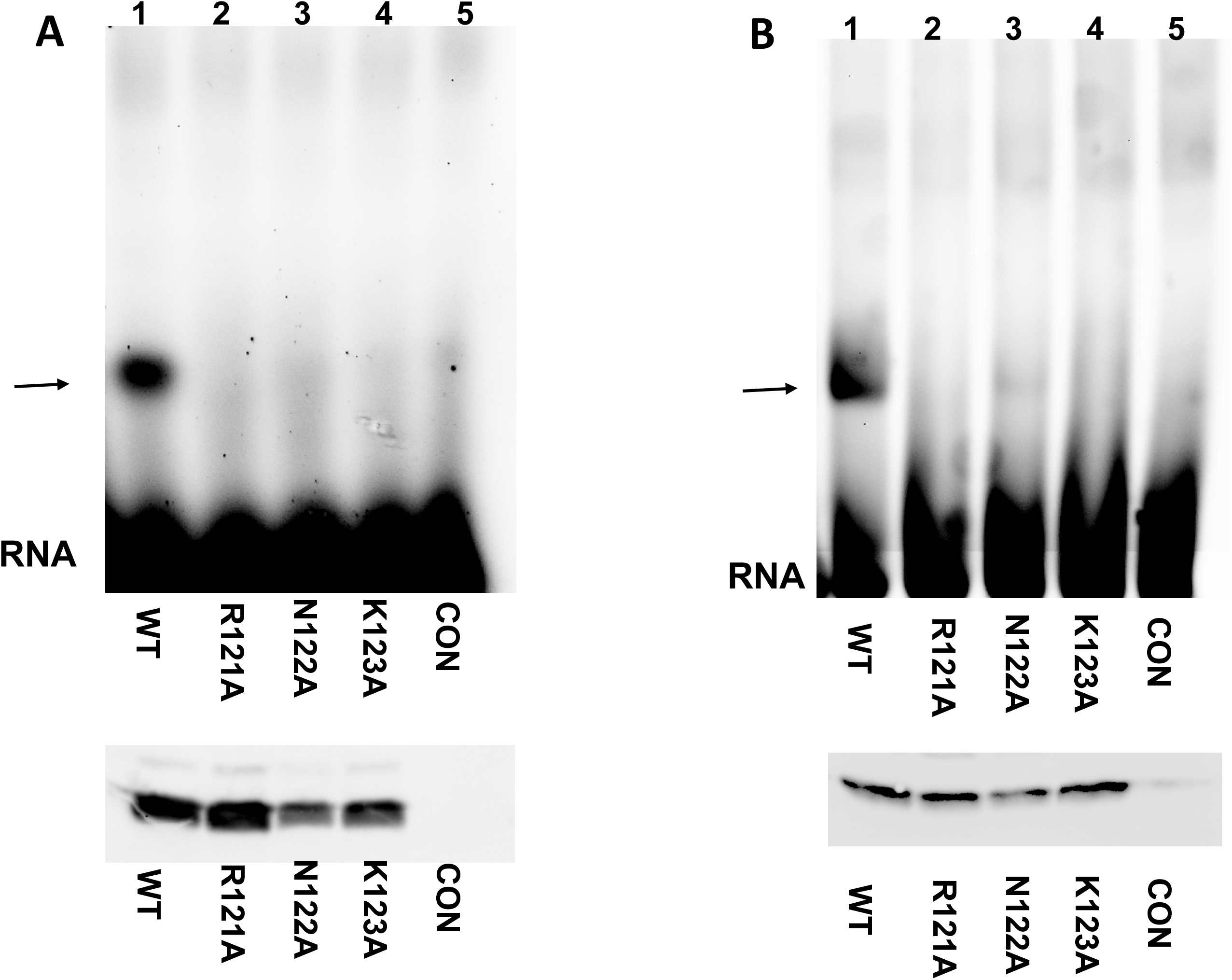
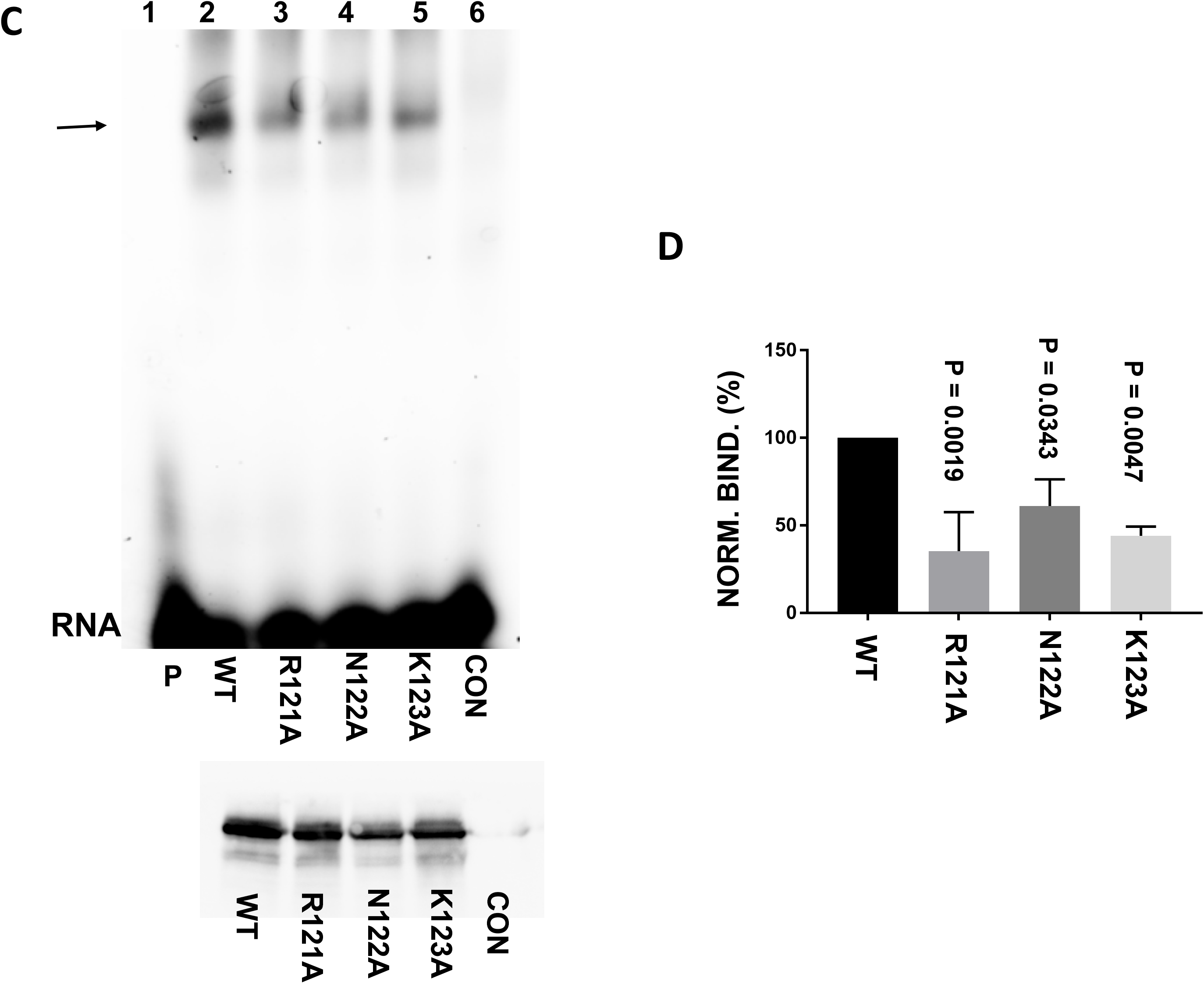
RNK epitope of La-domain is critical for binding 5’SL. A. Mutation of single amino acids of the RNK epitope abolishes binding of La-domain to 5’SL RNA. WT La-domain (lane 1) or the single amino acid mutants of RNK epitope (lanes 2-4) were expressed in HEK293 cells and cell lysates were analyzed for binding α2(I) 5’SL RNA by gel mobility shift. Lane 5, untransfected cells (CON). Bottom panel, expression of proteins in the same extract measured by western blot. B. The same experiment, but the binding to α1(I) 5’SL was analyzed. C. Mutation of individual amino acids of the RNK epitope reduces binding of full size LARP6. Experiment as in A, except the mutations were made in full size LARP6. D. Quantification of binding of the individual mutants of full size LARP6. Intensity of LARP6/5’SL RNA complex in gel shift experiment was normalized to expression of transfected LARP6 in the same extract and arbitrarily set as 100% for WT LARP6. N=3, error bars, +−1SD.

Because RRM increases the affinity of binding of La-domain we tested if the same mutations have a detrimental effect in the context of full size LARP6. When individual RNK mutations were tested in the context of full size LARP6 a significant decrease in binding to 5’SL was observed, but the binding was not completely abolished (Fig 5C). In three independent experiments, we normalized the signal of 5’SL/LARP6 complex from gel shift experiment to the expression of the protein in same extract determined by western blot and compared the normalized binding between the mutants. This analysis indicated that each individual mutation reduced the binding of full size LARP6 by about 50% (Fig 5D).

Next, we created double mutants in the context of full size LARP6; RK, RN or NK were changed into alanines, creating RK/AA, RN/AA and NK/AA mutants and repeated the analysis. Fig 6A and 6B show that the double mutations reduced LARP6 binding by 80-90%. To confirm this result, we made recombinant La-module with the RK/AA mutation and compared the binding to wt La-module. The binding of recombinant RK/AA mutant in the context of La-module was also drastically reduced (Fig 6C). The proteins used in this experiment are shown in the right panel of Fig. 6C. From these experiments we concluded that the RNK epitope is critical for sequence specific binding of LARP6 to 5’SL RNA. Because this epitope is found in a loop between the two secondary structures of La-domain, it is unlikely that alanine substitutions would cause gross structural alterations of the domain.

**Figure 6.**
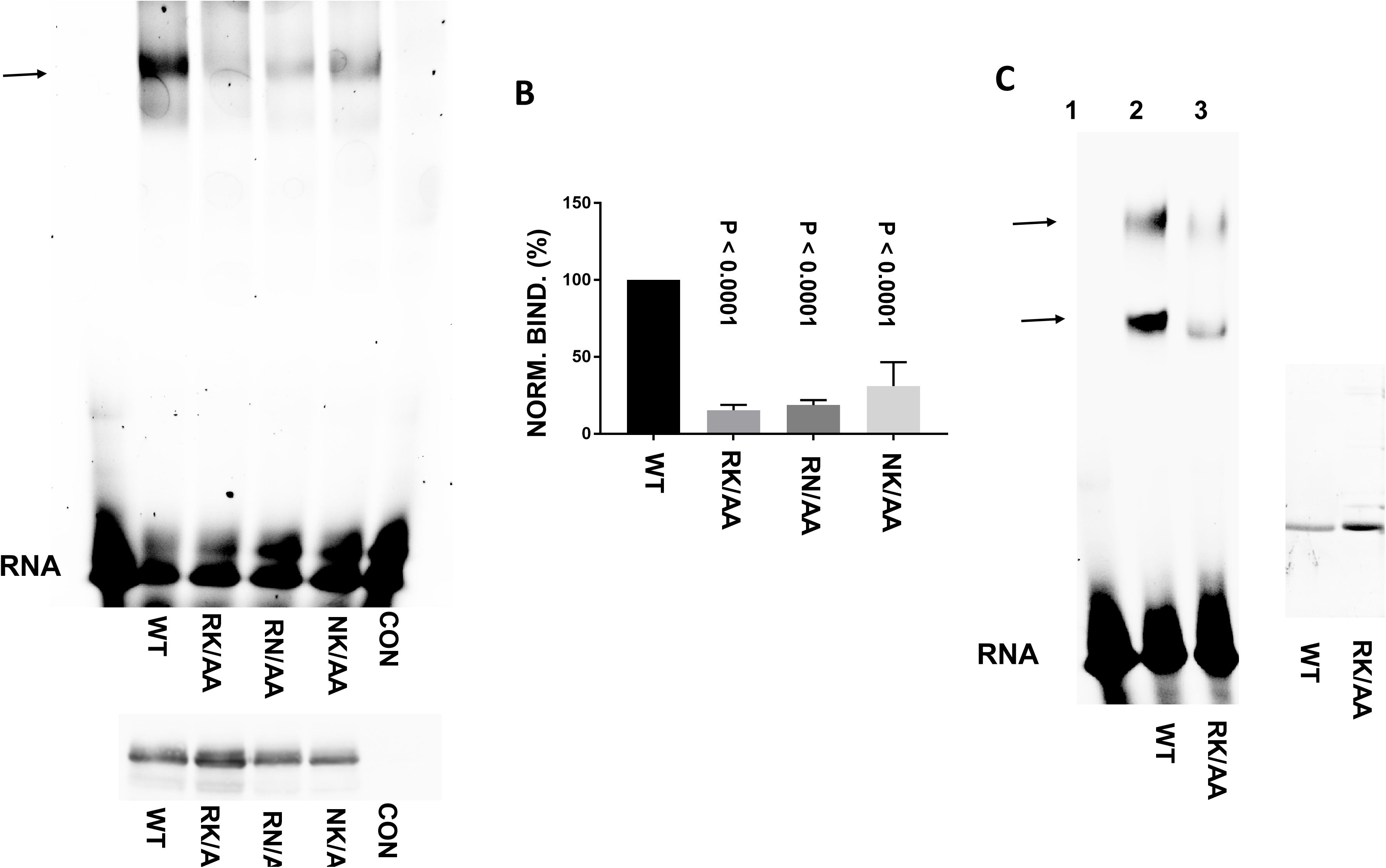
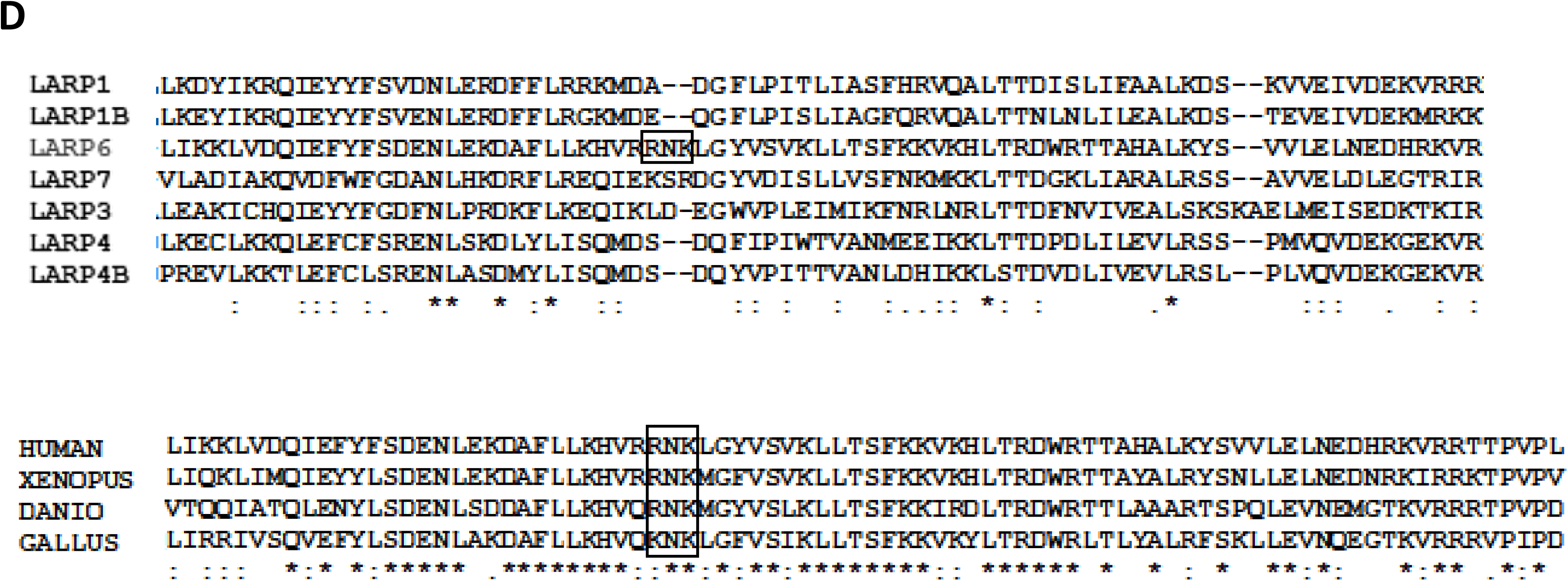
Double mutations of RNK epitope have dramatic effect on binding of full size LARP6. Two amino acids of RNK were simultaneously mutated as indicted (lanes 3-5) and the mutants were expressed in HEK293 cells and analyzed for binding α2(I) 5’SL RNA by gel mobility shift. Lane 1, RNA alone, lane 2, binding of wt LARP6, lane 6, non-transfected cells (CON). B. Quantification of binding of double mutants of full size LARP6 as in Fig. 5C. Binding of recombinant double mutant RK/AA. The double mutation was made in recombinant full size LARP6 and binding of pure protein (lane 3) was compared to that of wt LARP6 (lane 2). Lane 1, RNA alone. Right panel: recombinant proteins used in analysis. D. Sequence comparison of La-domains of human LARP family members (upper panel) and of La-domain of LARP6 of distant vertebrates (lower panel). RNK epitope is boxed.

### RNK epitope is not found in other LARPs, but is conserved in LARP6 of vertebrates

Other members of LARP superfamily have the La-module [4], but can not bind 5’SL. When the sequence of La-domains of human LARP superfamily members is compared, the RNK epitope is not found in other LARPs (Fig 6D). This explains why only LARP6 can bind 5’SL and is a compelling evidence that RNK epitope is not necessary for the folding of La-domain, but solely for interaction with 5’SL.

Because collagen 5’SL is found in collagen α1(I) and α2(I) mRNA of all vertebrates, LARP6 of all vertebrates must have the RNK epitope. Sequence comparison of LARP6 of distant vertebrates confirmed that the RNK epitope is conserved in vertebrates; only in birds the epitope is KNK instead of RNK (Fig 6D).

### The main site of crosslinking of La-module to 5’SL RNA maps to the asparagine of RNK epitope

To map the amino acid of LARP6 which forms covalent crosslink to 5’SL RNA upon UV irradiation we took advantage of the fact that the single amino acid mutants of RNK epitope bind 5’SL when analyzed in the context of full size LARP6 (Fig 5C). We surmised that a mutant that can bind 5’SL but can not be UV crosslinked has the amino acid involved in the crosslink formation changed. Therefore, we made R121A, N122A or K123A mutations in the context of La-module, prepared the recombinant proteins and compared their binding and crosslinking to α2(I) 5’SL RNA. In this experiment we adjusted the amount of mutant proteins to give similar binding in gel mobility shift experiment (Fig 7A), so we can directly compare their UV crosslinking efficiency (Fig 7B). While the R121A and K123A mutants crosslinked to 5’SL with the efficiency slightly less than wt La-module, the N122A mutant showed greatly reduced UV crosslinking signal (compare lanes 2 and 4). Thus, for similar binding activity, the UV induced attachment of N122A mutant to RNA was greatly reduced, strongly suggesting that the side chain of N122 is in close proximity to the RNA and forms the major crosslink. The fact that R121A and K123A mutants crosslinked less efficiently than wt La-module and that N122 mutation did not completely abolished the crosslinking, suggests that R122 and K123 participate in formation of minor crosslinks.

**Figure 7.**
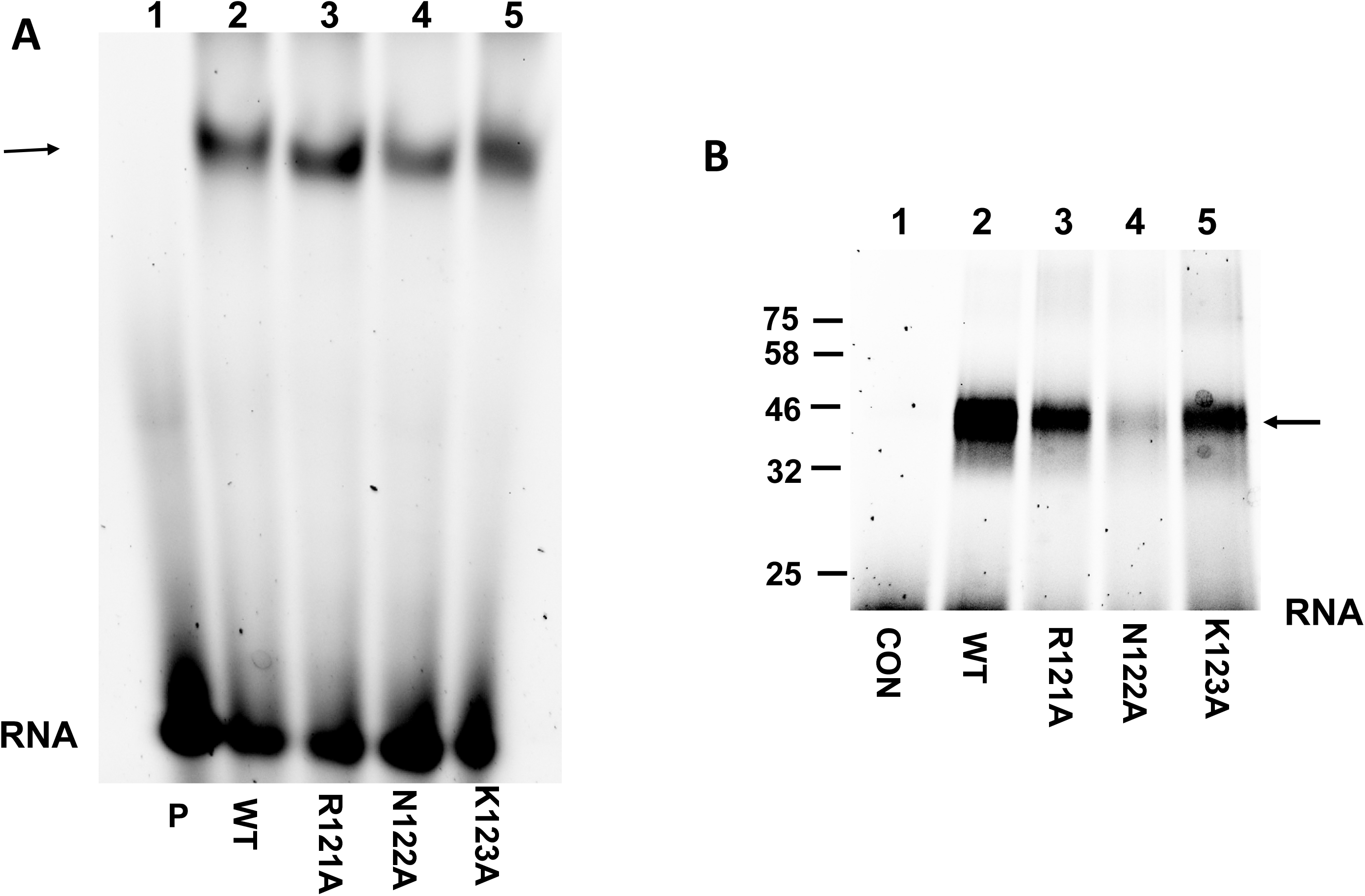
UV crosslinking of LARP6 to 5’SL RNA is by RNK epitope. A. Adjustment of binding activity of recombinant proteins. Gel mobility shift with the amount of recombinant wt La-module (lane 2) and the indicated mutants (lanes 3-5) adjusted to give similar binding signal to α2(I) 5’SL. RNA/protein complex is indicated by arrow. Lane 1, RNA alone (P). B. UV crosslinking of the same samples to α2(I) 5’SL. After UV irradiation the RNA/protein complexes were resolved on SDS-PAGE and visualized by imaging of RNA fluorescence. Lane 1, RNA without protein (CON). Arrow, RNA/protein complex.

### Mapping the interactions on 5’SL RNA

To assess which regions of 5’SL RNA are involved in binding of La-domain or La-module we performed RNA footprinting experiments. The 5’ end fluorescently labeled α1(I) 5’SL RNA or α2(I) 5’SL RNA were bound to recombinant La-domain or La-module and subjected to partial nuclease digestion. The regions occupied by the protein are then visualized as attenuated or absent cleavage products. In Fig 8A, α2(I) 5’SL RNA was bound by recombinant La-domain and the bound (lanes 4, 5 and 6) and free RNA (lanes 2, 3 and 7) was partially digested with benzonase. Benzonase is a nuclease which digests RNA irrespectively of the sequence and secondary structure [32]. As a marker, free RNA was partially digested with RNase A which preferentially cleaves after pyrimidines in single stranded regions (lane 1). This lane allowed us to assign the sizes of cleavage products. The cleavage products very close to the 5’end could not be clearly resolved, therefore, we could not determine the footprints at the ascending part of stem 1. Binding of La-domain protected the B1 portion of the bulge up to nucleotide 14 (indicated as FP1 in Fig 8A), it also protected nucleotides 32-34 just 5’ to the B2 portion of the bulge (FP2), and nucleotides close to the 3’ end, which correspond to the descending part of stem 1 (FP3). The location of these footprints on the RNA sequence is shown in Fig 8E.

**Figure 8.**
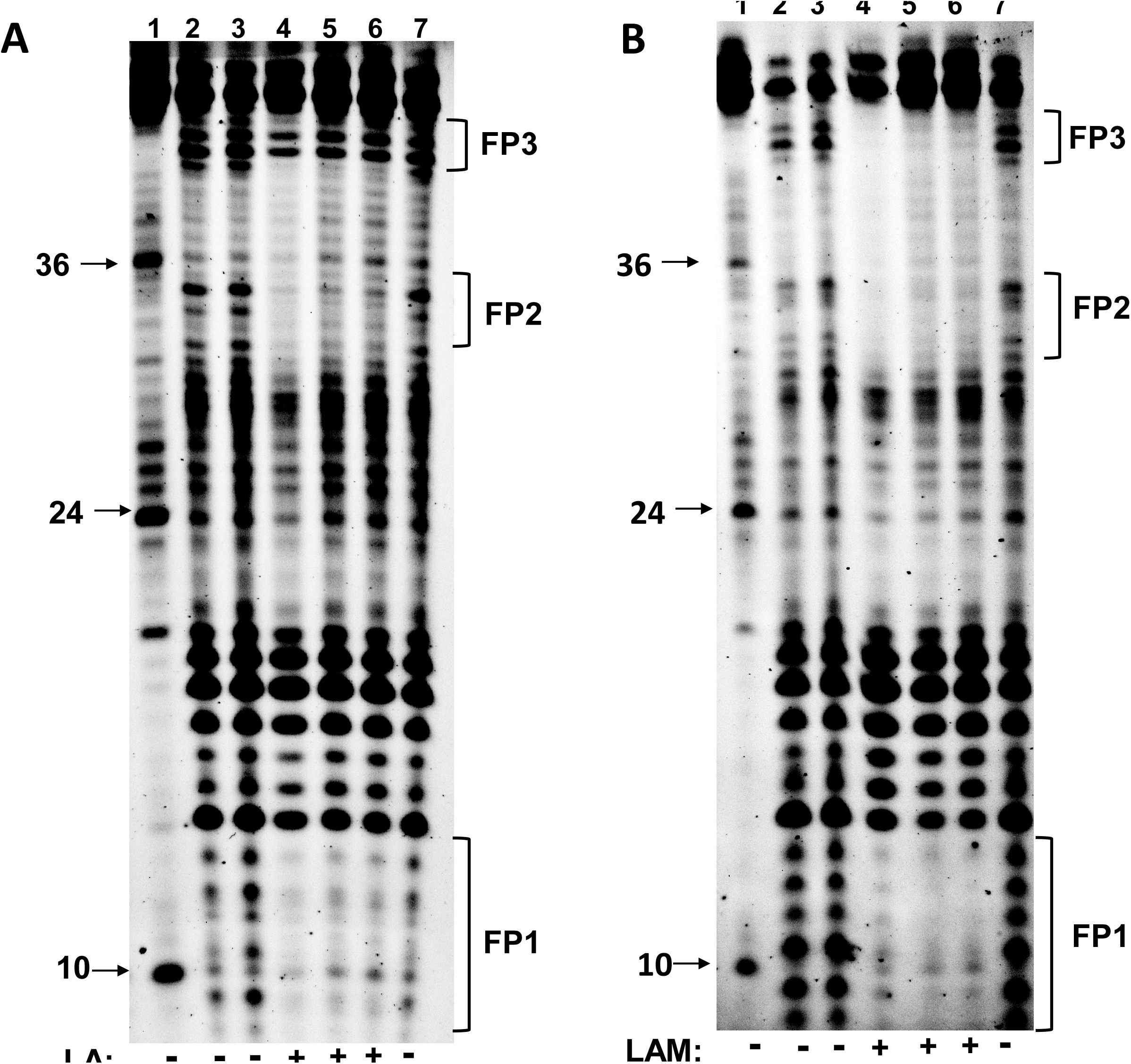
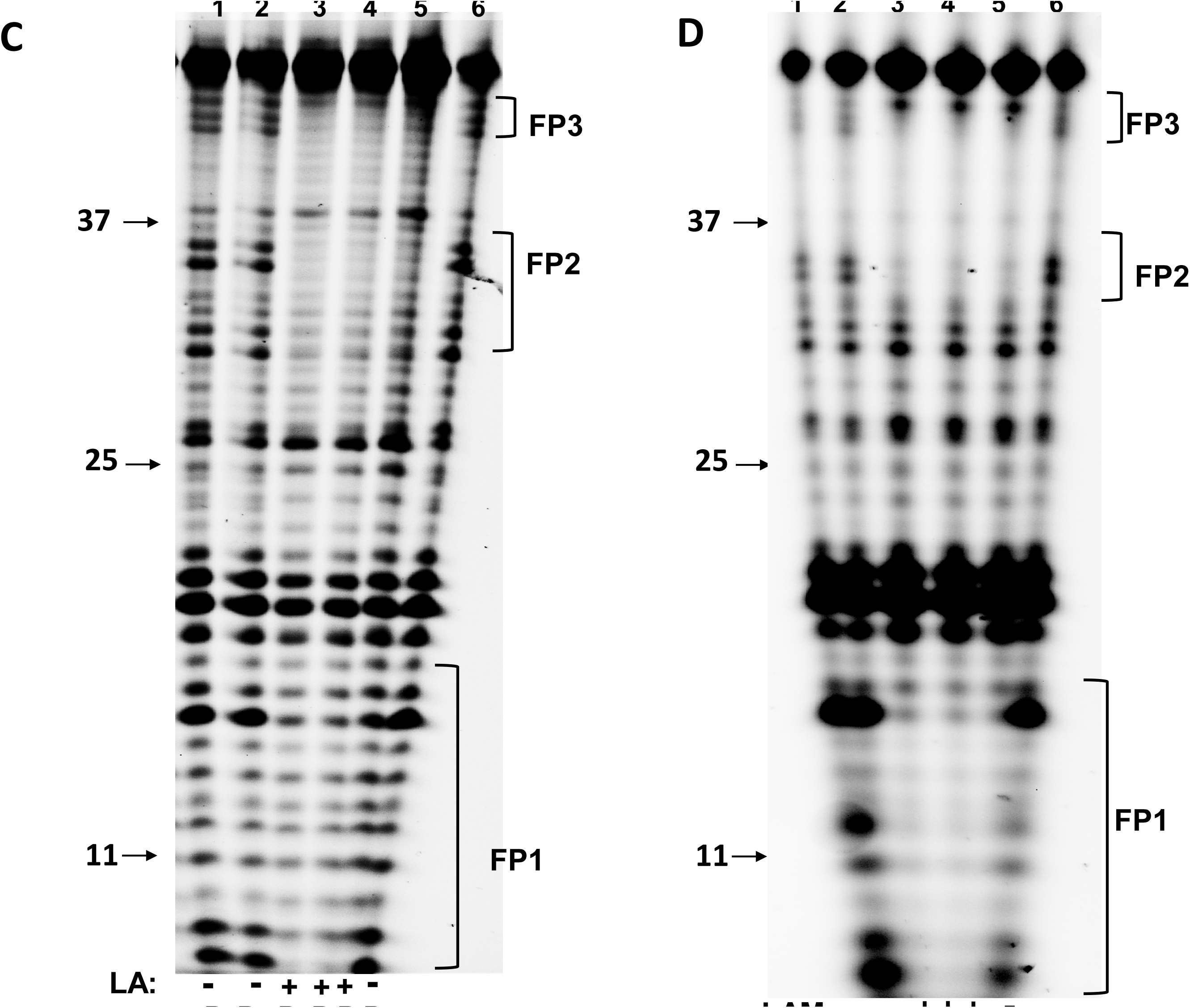
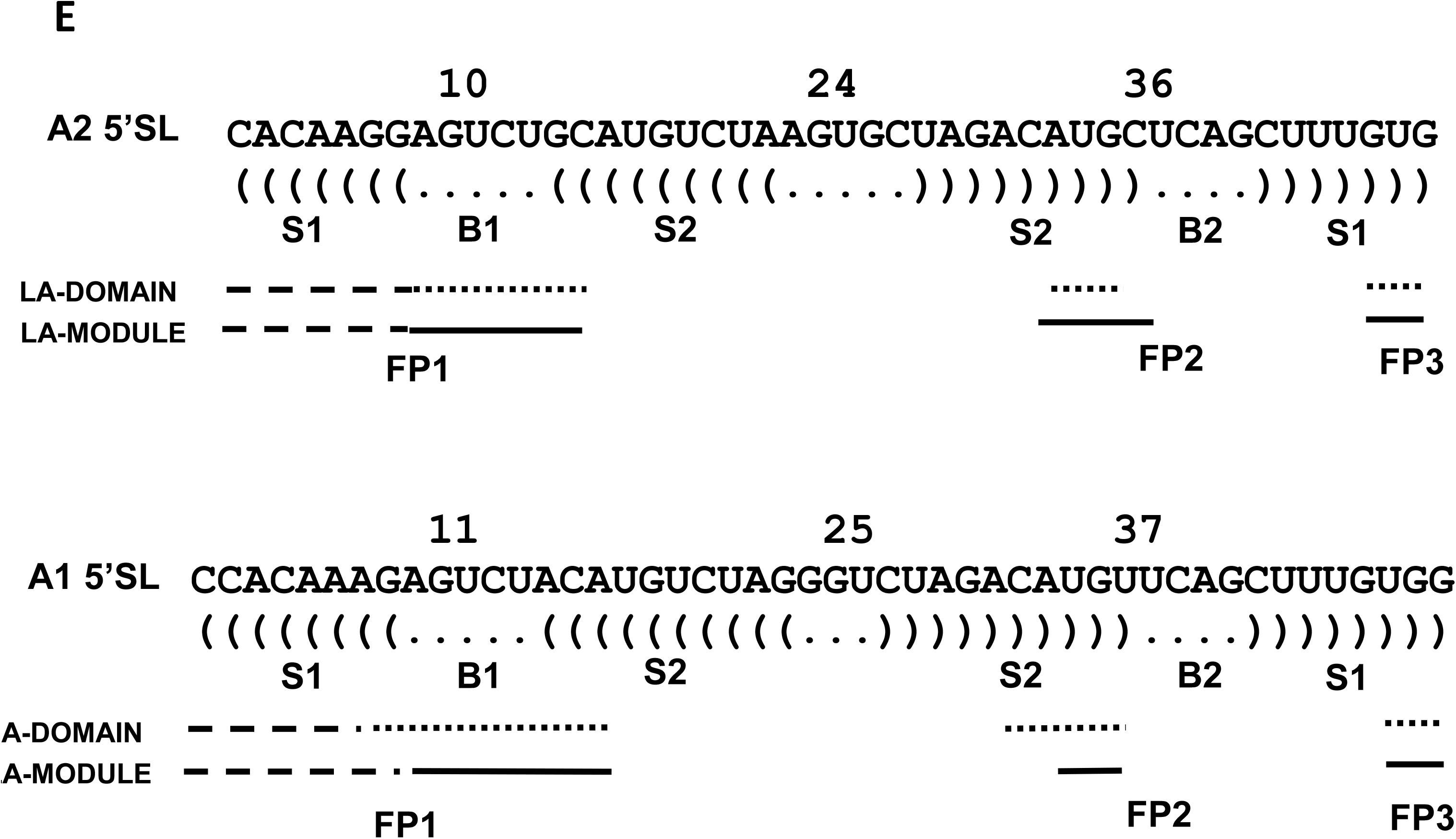
Footprints of La-domain and La-module on 5’SL RNA. A. Footprint of La-domain on α2(I) 5’SL RNA. 5’ end labeled 5’SL RNA, free (lanes 2, 3 and 7) or bound to recombinant La-domain (lanes 4-6) was partially digested with benzonase (B) and the fragments resolved on 8% denaturing acrylamide gel. The areas protected from nuclease cleavage are indicated as FP1, FP2 and FP3. Lane 1, partial digestion with RNase A (R), with the sizes of cleavage fragments indicated. B. The same experiment using recombinant La-module. C. Footprint of La-domain on α1(I) 5’SL. D. Footprint of La-module on α1(I) 5’SL. E. Schematic representation of footprints in the sequence of α2(I) 5’SL (upper panel) and α1(I) 5’SL (lower panel). Nucleotide numbers are on the top of the sequence and secondary structures are indicated under the sequence. Dotted line; footprints of La-domain, solid line; footprints of La-module, dashed line; region close to the 5’ end where footprints could not be determined. S1, lower stem, B1 and B2, bulge, S2, upper stem.

Binding of La-module (Fig 8B) showed similar protection of the B1 part of the bulge of α2(I) 5’SL up to the nucleotide 14 (FP1), and from nucleotides 31 to 35 adjacent to the B2 portion of the bulge (FP2). A protection close to the 3’ end is also seen (FP3). All footprints were stronger with La-module than the footprints seen with La-domain, but the extent of the protected regions was not significantly increased. This suggested that RRM increases the strength of binding, but does not protect additional regions of 5’SL RNA (Fig 8E).

Footprinting of α1(I) 5’SL RNA by La-domain is shown in Fig 8C. There was weak protection of the B1 segment of the bulge (FP1) up to the nucleotide 16, protection of nucleotides 33-35 adjacent to the B2 segment (FP2) and protection of nucleotides close to the 3’ end, which correspond to the descending part of stem 1 (FP3). This pattern was similar to that seen with the α2(I) 5’SL (Fig 8E).

Fig 8D shows the protection of α1(I) 5’SL RNA by La-module. La-module strongly protected the B1 segment of the bulge and part of stem 2 up to the nucleotide 16 (FP1). La-module also showed strong protection of nucleotides 32-34, which are adjacent to the B2 segment of the bulge (FP2) and of nucleotides close to the 3’ end, which correspond to the descending part of stem 1 (FP3). Again, the footprints were stronger, but not significantly larger than with La-domain (Fig 8E).

Overall, these results indicate that the footprints on α1(I) and α2(I) 5’SL cover the equivalent parts of the RNA, they are stronger when RRM is present (La-module) but the area covered by the footprints is not enlarged compared to footprints with La-domain (Fig 6E). This is consistent with the notion that La-domain primarily contacts the RNA and that RRM increases the affinity of binding.

### Mapping of UV crosslinking site on 5’SL RNA

Fig 3D shows that La-module can be UV cross-linked to 5’SL RNA more efficiently than La-domain, allowing us to map the site of crosslinking on 5’SL RNA using La-module. To increase the crosslinking efficiency we synthesized α1(I) 5’SL RNA by in vitro transcription with 50% uridines substituted with 4-thio uridines [33] and radiolabeled the RNA at the 5’end. This RNA was longer than the 5’SL RNAs used in footprinting experiments and is shown in Fig 9A. After UV crosslinking of La-module to this RNA, the crosslink was purified by SDS-PAGE electrophoresis and subjected to partial NaOH hydrolysis. The hydrolysis produces a series of RNA fragments of increasing size until the position of crosslink is reached. After the crosslink the RNA fragments contain covalently attached protein and have greatly decreased electrophoretic mobility and accumulate at the top of the gel. Thus, the crosslinked site can be mapped as a discontinuity in NaOH generated ladder [34]. Fig 9A, shows NaOH cleavage products of RNA without UV crosslinking (lane 1) and of the RNA crosslinked to La-module (lane 2). The ladder showed the discontinuity after nucleotide 67 (arrow), what mapped to the B2 portion of the bulge (Fig 9A).

**Figure 9.**
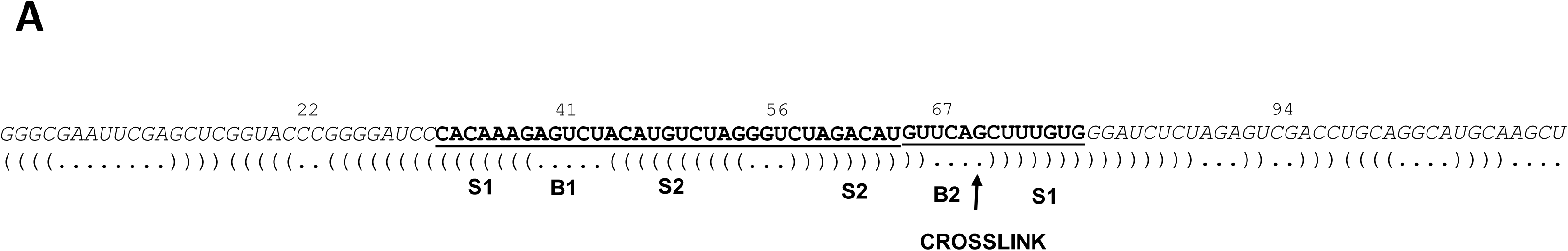
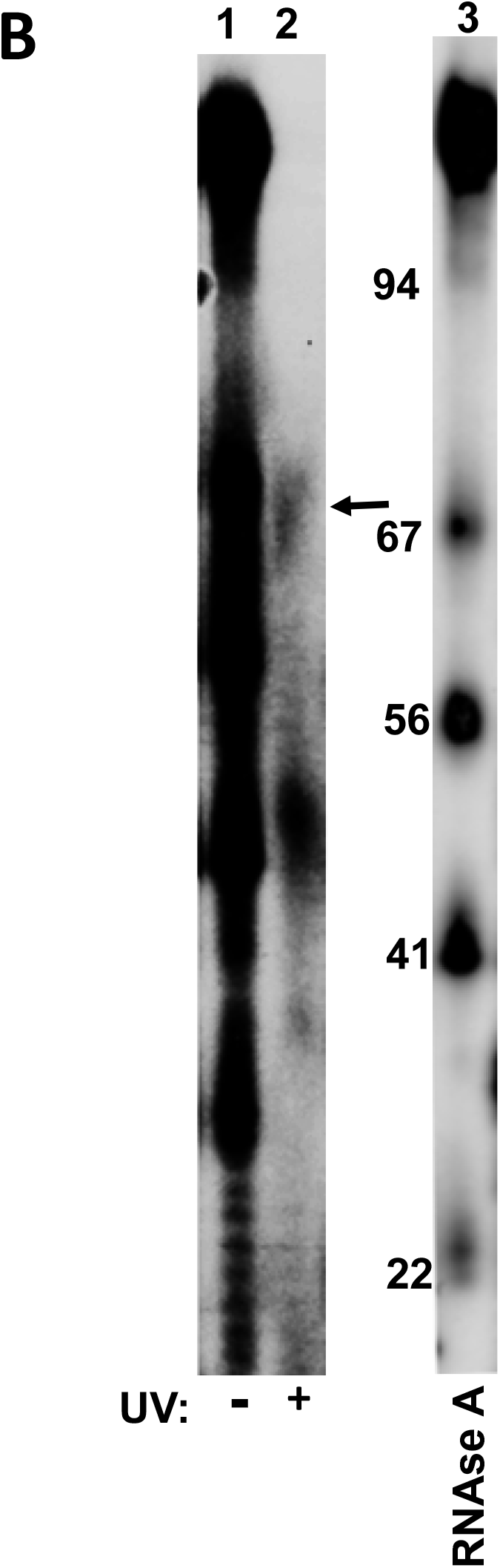
Mapping of crosslinking site on 5’SL RNA. A. Sequence of RNA used for crosslinking. Nucleotide numbers are on the top of the sequence and secondary structure is indicated under the sequence. Underlined is the sequence of α1(I) 5’SL with S1, lower stem, B1 and B2, bulge and S2, upper stem. Position of crosslink is indicated by arrow. B. Mapping of crosslink. Free RNA (lane 1) or RNA with La-module covalently crosslinked (lane 2) or were partially hydrolyzed by alkali and fragments resolved on 8% PAGE/urea gel. Arrow indicates the discontinuity of ladder. Lane 3, same RNA partially digested with RNase A with sizes of fragments indicated. This lane was run on the same gel as lanes 1 and 2, but exposed shorter.

## Discussion

LARP6 is the specific regulator of translation of type I collagen mRNAs and recognition of 5’SL of collagen mRNAs by LARP6 is critical for fibrosis development [27, 28]. Therefore, LARP6 binding inhibitors show promise as antifibrotic compounds [28], because they overcome the main drawback of antifibrotic drugs currently in development; the lack of specificity. This work provides insight into the molecular recognition between LARP6 and 5’SL, as the first step to help rational design of such inhibitors. The main findings are: 1. La-domain of LARP6 is necessary and sufficient to bind 5’SL in sequence specific manner (Figs 1 and 2), 2. RRM domain serves to stabilize the interaction of La-domain with 5’SL RNA (Fig 3), but does not show extensive contacts with the RNA (Fig 8). 3. The short epitope of La domain, RNK, is critical for binding (Figs 5 and 6), suggesting that limited number of amino-acids convey the sequence specific recognition. 4. The direct contact between LARP6 and 5’SL RNA is between N122 of the RNK epitope and one of the nucleotides within the B2 portion of the bulge (Figs 7 and 9). 5. The sequences at the C-terminus of La domain, which form a linker with RRM, also participate in binding, but not in the sequence specific manner. Based on the NMR structure of La-domain, the loop containing RNK epitope and this linker form a pocket with flexibile structure. This work suggests that 5’SL RNA binds this pocket.

It has been reported previously that La-domain alone can not bind 5’SL [11]. However, that work used isothermal titration calorimetry (ITC) to measure the binding, while we employed gel mobility shift to visualize the RNA/protein complexes. The difference in detection by these methods may be the cause for the discrepancy. However, during the course of our experiments we noted that the binding activity of La-domain rapidly deteriorates upon manipulations and storage of the protein, what is not the case with La-module. Therefore, in our experiments we always used fresh samples of La-domain. With this caveat, we were able to show that the recombinant La-domain or La-domain expressed in cells can bind 5’SL RNA in sequence specific manner and can discriminated between the single nucleotide mutants of 5’SL (Fig 2).

The identification of RNK motif as critical sequence for recognition of 5’SL RNA is an important new finding. All LARP family members have the highly conserved La-domain, but their La-domain lacks the RNK sequence, what can explain why they can not bind collagen 5’SL. Mutations of individual RNK amino acids completely abolish the interaction of La-domain with 5’SL RNA. In the context of full size LARP6 the individual mutations are less detrimental and reduce binding by ~50%. However, simultaneous mutations of two amino acids of the epitope reduce binding by ~90% (Fig 5), indicating the contribution of all three amino acids to binding. The RNK epitope resides in a loop that connects the second α-helix to the first β-sheet strand of La-domain. Thus, the epitope is not part of any secondary structure and is exposed on the surface of the protein. In the NMR spectra of La-domain the resonances of atoms in this loop could not be assigned [11], suggesting an inherently flexible structure.

Besides the RNK epitope, La-domain requires amino acids at the C-terminus, which connect La-domain to RRM. Truncation of 8 amino acids at the C-terminus is detrimental to La-domain binding, but the binding can be restored by substituting random amino acids. We have found one amino acid sequence which increased the binding (Fig 4B), what suggested that the sequences appended to the C-terminus can modulate the strength of interaction of the La-domain. Previous work indicated that this linker is important in binding of La-module, as well [11]. The linker also has a flexible structure and together with the RNK loop forms a highly dynamic cavity (PDP ID: 2MTF). It can be speculated that the dynamics of this pocket is needed to establishing the contacts with the bulge of 5’SL RNA.

The crystal structure of La-module of LARP3 [35] and LARP7 [36] bound to short poly-U RNA have been solved. In both cases the RNA is sandwiched in the binding pocket between La-domain and RRM with both domains making contacts with the RNA. Our results indicate that the recognition of 5’SL RNA by LARP6 is different and that it involves a pocket formed by the sequences of La-domain.

When La-domain and La-module are titrated with 5’SL RNA separately both proteins bind the RNA with similar affinity (Fig 3A). However, when they compete for the RNA, La-module outcompeted binding to La-domain when present at 1.5-fold higher molar ratio, while La-domain at 14-fold higher molar ratio had no effect on the La-module binding (Fig 3B and 3C). This suggests that the complexes formed with La-module are more stable and that the role of RRM is to increase the stability of the 5’SL/LARP6 complex. The area of 5’SL RNA that is protected from nuclease digestion by La-domain includes the bulge and stem 1, however, this area is not significantly larger when La-module is bound, just the protection is stronger (Fig 8). This supports the conclusion that RRM does not make extensive additional contacts with the RNA. RRM alone can not bind 5’SL RNA and in trans can not augment the binding of La-domain (Fig 1), therefore, it has to be tethered to 5’SL by the La-domain. How RRM increases the affinity of La-domain for 5’SL RNA can not be discerned from this study and must await determination of the structure of La-module.

For now, only a putative model can be proposed that the initial sequence specific contact is established between the RNK epitope of La-domain and the nucleotides of the bulge. The close proximity of the N122 and the nucleotides at B2 side of the bulge (Figs 7 and 9) has been suggested by UV crosslinking, The weaker crosslinking of the other two amino acids of the epitope does not necessarily mean that they are not in close proximity, but that their side chains may be less reactive to the UV light or contacting less UV reactive nucleotides. Nevertheless, the results clearly point to the importance of these amino acids in sequence specific recognition of 5’SL. After the initial sequence specific recognition contact is established, what may be facilitated by the inherent flexibility of the binding pocket, the RRM domain closes the conformation to increase the half life of the complex.

It has always been surmised that protein-RNA binding involves large surface area and that it can not be dissociated by small molecules [37]. This may be true for LARP6 that is already bound to 5’SL because of high affinity of binding, however, the involvement of the relatively small RNK epitope in the initial recognition suggests that blocking the RNK docking to 5’SL by small molecules would render LARP6 inactive in collagen biosynthesis. Finding such inhibitors is of utmost importance, because RNK epitope is not found in other LARPs and 5’SL RNA is found only in type I and type III collagen mRNAs, thus, such inhibitors may be specific for type I collagen expression in fibrosis.

## Material and methods

### Preparation of recombinant proteins

For expression of La-domain of human LARP6 in E. coli the sequence of human LARP6 cDNA encoding for amino acids 81-182 was cloned in pET15b vector, while for expression of RRM domain the sequence encoding amino acids 183 to 298 was cloned. The construct expressing recombinant La-module was described in [10]. Site directed mutagenesis was performed with QuickChange II kit (Stratagene) according to manufacturer instructions and the identity of mutants was verified by sequencing. pET15b vectors were transformed into Rosetta cells and protein expression was induced with 1mM IPTG for 3h. Cells were sonicated and recombinant proteins purified in a single step on Ni-agarose column according to manufacturer instructions (Invitrogen). Protein samples were dialyzed against PBS with 5 mM MgCl_2_ and 0.1% glycerol for 2h and used immediately in experiments. The purity of preparations was analyzed by SDS-PAGE and Coomasie staining.

### Expression in mammalian cells

La-domain and RRM described above were cloned into pCDNA3 vector containing 2HA tags at the N-terminus for La-domain and 2HA tags at the C-terminus for RRM domain. Full size LARP6 construct was described in [10]. Site directed mutagenesis was performed with QuickChange II kit (Stratagene) according to manufacturer instructions and the identity of mutants was verified by sequencing. The constructs were transfected into HEK293 cells and two days after transfection cells were lysed in PBS containing 5 mM MgCl_2_, 0.5% NP-40. 0.5% glycerol and protease inhibitors and cell extracts were used in gel mobility shift experiments and western blots immediately after preparation.

### Gel mobility shift experiments

The binding reactions contained 5-40 μL of recombinant proteins or HEK293 cell extract, 20 nM of 5’SL RNA probe fluorescently labeled at the 5’ end (Dharmacon) and 10 μg of yeast t-RNA in total volume of 50 μl. The probe RNAs had sequence of human collagen α1(I) and α2(I) 5’SL and were 48 and 46 nucleotides long, respectively. Reactions were incubated at RT for 15 min and resolved on 6% native acrylamide gels, as described [10]. For cross-competition experiments recombinant proteins were mixed in the indicated amounts prior to addition of 5’SL RNA probe. Imaging of the gels was done by Biorad ChemiDoc MP imaging system and intensity of bands was quantified by Image J program.

### Western blots

The same cell extracts used in gel mobility shift experiments were analyzed by western blots. 30 μl of extract was resolved on 10 or 12% SDS-PAGE gels and after transfer the membranes were probed with 1:1000 dilution of anti-HA antibody (Sigma). Imaging of blots was done by Biorad ChemiDoc, 2.3.0.07 imagining system and intensity of bands was quantified by Image J program.

### Fluorescence polarization

Fluorescence polarization (FP) was measured as described. Recombinant La-domain and La-module at concentration of 3.8 μM were titrated with increasing amounts of fluorescently labeled α2(I) 5’SL RNA in PBS, 5mM MgCl_2_, in total volume of 25 μl. FP was measured in 384-well plates using Biotek Synergy H1 plate reader. FP of blanks without protein was subtracted from these readings to obtain the protein dependent FP values.

### UV crosslinking

For analysis of crosslinks the reactions used for gel mobility shift were irradiated with UV light of 254 nM at distance of 5 cm for 15 min. After crosslinking, samples were resolved on 10 or 12% SDS-PAGE gels and the gels were imaged as above. For preparative crosslinking, ds-oligonucleotide containing the sequence of human collagen α1(I) 5’SL was cloned into BamHI site of pGEM3 vector, the construct was linearized by HindIII and used as template for in vitro transcription. The in vitro transcription reaction was supplemented with 4-thio uridines (Sienna Bioresearch) to incorporate 4-thio uridines at 50% of total uridines and the RNA was 5’ end labeled with ^32^P. Crosslinking was performed with 30 μM recombinant La-module in 200 uL reaction and resolved on 10% preparative SDS-PAGE gel. After radiography, the crosslinked band was excised, eluted and subjected to partial NaOH hydrolysis. 50 μl of sample was hydrolyzed with 25 μl of 50 mM bicarbonate buffer, pH 9.2 for 4 min at 95C and fragments were resolved on 8% sequencing gel and visualized by autoradiography. As a marker, the same RNA was partially digested with RNAse A and digestion products were run on the same gel.

### RNA footprinting

The 5’SL RNAs used in gel mobility shift experiments and labeled at the 5’ end by fluorescein were used free or bound to recombinant La-domain or La-module and were digested in 50 μl PBS, 5 mM MgCl2 with 2.5U of Benzonase (AMD Millipore) for 6 min at RT, phenol/chlorophorm extracted and analyzed on 8% acrylamide/8M urea gel. For RNase A generated ladder, free RNAs were digested with 100 ng of RNAse A for 8 min at RT and processed as above. The gels were imaged with Biorad ChemiDoc MP imaging system.

### Immunostaining

pCDNA3 vectors expressing HA-tagged full size LARP6 or La-domain were transfected into HEK293 cells and 24h after transfection the cells were fixed and incubated with anti-HA primary antibody (Sigma), followed by incubation with fluorescent secondary antibody. Non-transfected cells served as negative control. The images were taken with Keyence BZ-X710 microscope at 60x magnification.

