## Supporting Information for "Characterization of sequence specific binding of LARP6 to the 5’ stem-loop of type I collagen mRNAs and implications for rational design of antifibrotic drugs"

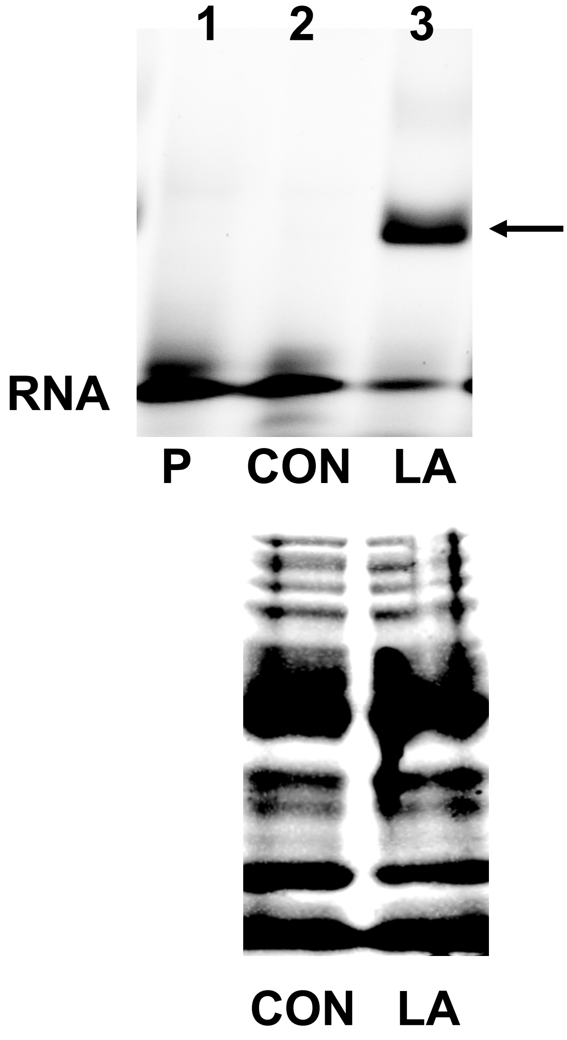


Supplemental figure 1. Binding to 5'SL in whole bacterial lysate. Upper panel: gel mobility shift with whole bacterial lysate without expressing La-domain (lane 2) or with expressing La-domain (lane 3). Lane 1 is 5'SL RNA probe alone. Bottom panel: proteins in whole bacterial lysates used in the gel mobility shift stained with Coomassie blue.

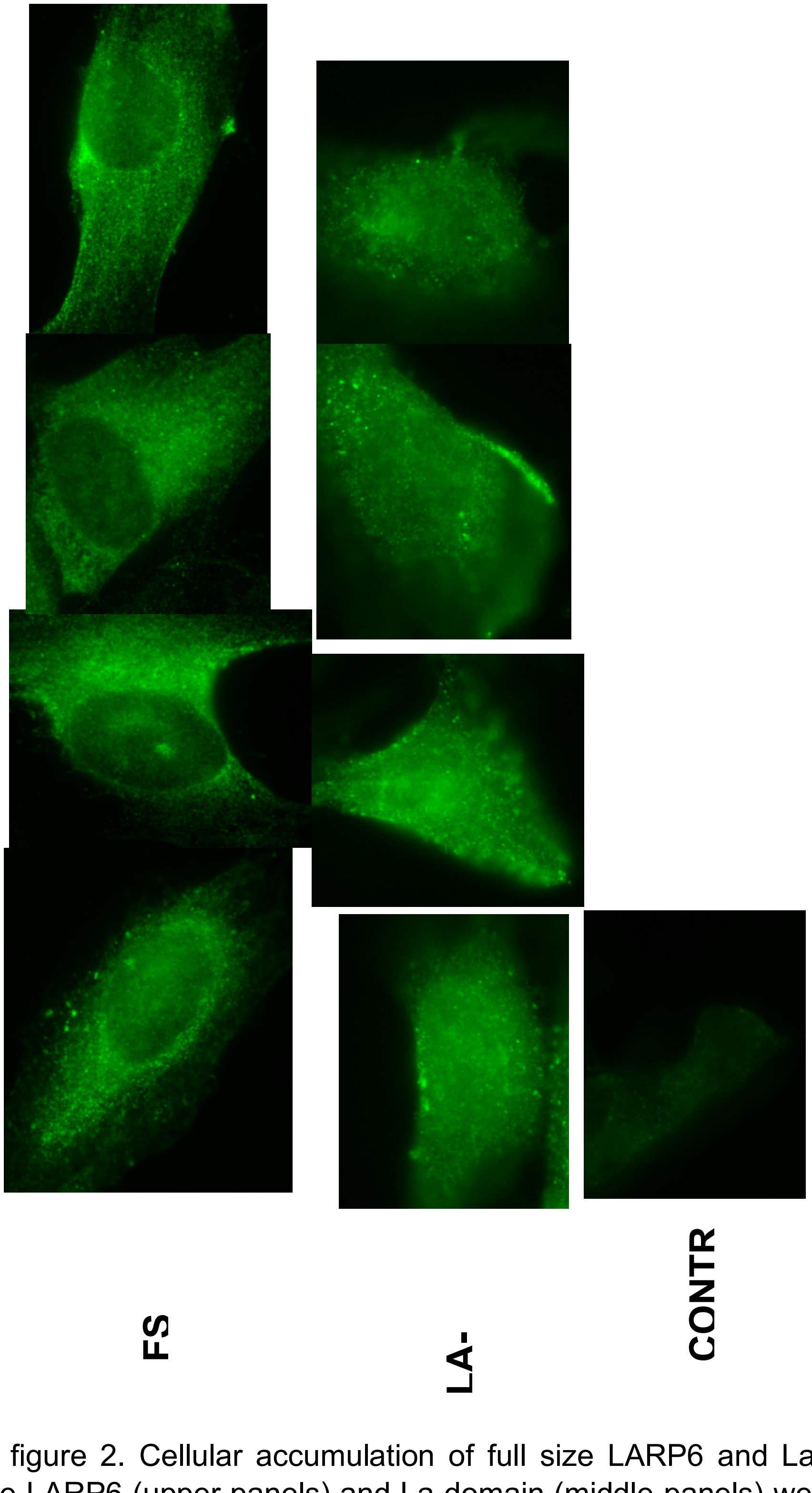


Supplemental figure 2. Cellular accumulation of full size LARP6 and La-domain. HA-tagged full size LARP6 (upper panels) and La-domain (middle panels) were transfected into HEK293 cells and immunostained with anti-HA antibody. Bottom panel: control untransfected cells.
